# Keystone engineering enables collective range expansion in microbial communities

**DOI:** 10.1101/2025.01.11.632568

**Authors:** Emrah Şimşek, César A. Villalobos, Kinshuk Sahu, Zhengqing Zhou, Nan Luo, Dongheon Lee, Helena R. Ma, Deverick J. Anderson, Charlotte T. Lee, Lingchong You

## Abstract

Keystone engineers profoundly influence microbial communities by altering their shared environment, often by modifying key resources. Here, we show that in an antibiotic-treated microbial community, bacterial spread is controlled by keystone engineering affecting dispersal—an effect hidden in well-mixed environments. Focusing on two pathogens, non-motile *Klebsiella pneumoniae* and motile *Pseudomonas aeruginosa*, we found that both tolerate a β-lactam antibiotic, with *Pseudomonas* being more resilient and dominating in well-mixed cultures. During range expansion, however, the antibiotic inhibits *Pseudomonas*’ ability to spread unless it is near *Klebsiella*—*Klebsiella* degrades the antibiotic to create a “clear zone” that allows *Pseudomonas* to expand, at the expense of *Klebsiella*’s own growth, thus acting as a keystone engineer. As *Pseudomonas* spreads, it competitively suppresses *Klebsiella*. Our modeling and experimental analyses reveal that this keystone effect operates at a millimeter scale. We also observed similar keystone engineering by a *Bacillus* species isolated from a hospital sink, in both pairwise and eight-member bacterial communities with its co-isolates. These findings suggest that spatially explicit experiments are essential to understand certain keystone engineering mechanisms and have implications for surface-associated microbial communities like biofilms, as well as for diagnosing and treating polymicrobial infections involving drug-degrading, non-motile (e.g., *Klebsiella*), and drug-tolerant, motile (e.g., *Pseudomonas*) bacteria.

## Introduction

A keystone species has a disproportionately large impact on its community relative to its abundance^1,2^. For example, removing the sea star *Pisaster ochraceous* drastically reduces biodiversity in its ecosystem because it allows mussel, its prey, to overgrow^1^. An ecosystem engineer affects other species in its community by modifying their environment. For example, beavers shape freshwater habitats that benefit many other species^3,4^. A species that meets both definitions is called a *keystone engineer*^5^.

In microbial communities, keystone engineers are critical for maintaining biodiversity^6,7^ and community functioning^8–13^, including their impact on human health^14–17^. The keystone species concept originated in natural communities, but the inherent complexity of the latter makes it challenging to uncover their underlying ecological mechanisms. To alleviate this challenge, simplified microbial communities with two to several members have been assembled to extract fundamental mechanistic insights, which can then be tested in increasingly complex communities^18^. Examples include predicting large community outcomes from pairwise interactions^19^, revealing limitations of pairwise information for larger communities^20^, demonstrating how interspecies interactions during chemotaxis and growth shape marine community assembly^21^, showing bioelectric signals mediates interspecies attractive interactions^22^, and illustrating how transient invaders can trigger state switching in a two-member community through pH modification^23^. This approach has also revealed keystone roles for bacteria in shaping community composition^7,24^, stability^25^, biomass production^12,13^, biofilm formation^26^, pathogen resistance^25^, and antimicrobial tolerance^26,27^.

Microbial keystone engineers often play their role by breaking down complex resources into products utilizable by other microbes^14,17,24,28–31^. This role can also entail changing the environmental pH^12,24^ or modulating the host immune response^15,16,32,33^. For instance, despite its low abundance, *Porphyromonas gingivalis* acts as a keystone that promotes overgrowth of the oral microbiota, which can cause inflammation^16^.

Whether a microbe is a keystone engineer depends on context. For instance, in a synthetic mouse gut microbiota, *Bacteroides caecimuris* serves as a keystone engineer only when the polysaccharide inulin is the carbohydrate source^24^, as *B*. *caecimuris* inhibits the growth of other species by acidifying the culture through inulin fermentation^24^. Likewise, *Bacteroides uniformis* can benefit or compete with butyrate-producing gut bacteria depending on the available carbohydrate sources^31^.

While previous studies on keystone engineering have focused predominantly on growth or community structure^14–17,24,28–33^, spatial structure is a ubiquitous environmental context for microbes. Many microbes live in spatially structured communities, such as biofilms^34–36^. In such an environment, survival depends on both mobility and effectiveness in nutrient utilization^37–40^. The mobility can be intrinsic or through interactions with other community members. For example, *Escherichia coli* can be transported by *Paenibacillus vortex*^41^, and *Pseudomonas aeruginosa* variants hitchhike on shared biosurfactants^42,43^. Biosurfactant production enables collective spatial expansion (called swarming) but comes at a significant growth cost, and cheater variants that do not produce biosurfactants can destabilize this strategy during a prolonged spatial expansion of a population^43^. When invading a spatial environment, *Pseudomonas’* biosurfactants can enable the displacement of competing *Klebsiella*, which previously colonized the environment^44^. These examples underscore the critical need for spatially explicit experiments to uncover microbial interactions. However, due to the lack of such well-controlled experiments, keystone engineering in the context of spatial expansion has not been demonstrated before.

Here, combining mathematical modeling and experiments, we reveal that in a microbial community treated with a β-lactam antibiotic, an immotile species acts as a keystone engineer enabling the spatial expansion of a motile species when sufficiently close. The immotile species detoxifies the environment by degrading the antibiotic, which would otherwise suppress the dispersal—but not the growth—of the motile species. This enables the motile species to expand, and subsequently suppress the growth of the immotile species through an apparent competition, reducing it to a minority. These dynamics constitute keystone engineering by the immotile species, which has a disproportionately large driving role on the spatial expansion of its community compared to its extremely low final relative abundance.

We demonstrated this keystone engineering in two pairwise communities: 1) clinical isolates of β-lactamase (Bla)-producing *Klebsiella pneumoniae* (*Klebsiella*) as the keystone engineer and *Pseudomonas aeruginosa* (*Pseudomonas*) as the motile species under cefotaxime (CTX) treatment, and 2) a nonpathogenic *Bacillus paranthracis* (Bp) with Bla genes (Supplementary Table 1) as the keystone engineer and another, motile *Pseudomonas* strain (Pa22), co-isolated from the same hospital sink, under carbenicillin (CB) treatment. Finally, we demonstrated the applicability of Bp’s keystone engineering on Pa22 in a synthetic community with six additional isolates from the same sink sample.

Based on our results, we hypothesize that a pathogen could appear responsible for a β-lactam resistant infection even though it does not encode Bla or degrade β-lactams (e.g., *Pseudomonas*), potentially leading to misleading diagnoses. Furthermore, our findings have broader implications for biofilm growth in mixed communities of pathogenic and nonpathogenic microbes in natural environments, such as hospital sinks.

## Results

### *Klebsiella* enabled *Pseudomonas* range expansion under β-lactam treatment at its own expense

We tested the effect of a β-lactam antibiotic (CTX) treatment on the growth (i.e., biomass production) of *Pseudomonas aeruginosa* PA14 (*Pseudomonas*) and a Bla-producing clinical isolate of *Klebsiella pneumoniae* (*Klebsiella*)^45^ in well-mixed liquid cultures. *Pseudomonas* and *Klebsiella* are prevalent pathogens in many common infection types^46,47^ (Supplementary Fig. 1), can co-colonize patients^48^, and co-occur at human infection sites^49^ or urinary catheters^50,51^, suggesting that they have a strong niche overlap, making their community experiments clinically and ecologically relevant. We first grew *Klebsiella* and *Pseudomonas* in well-mixed pure cultures without or with CTX (2.5 𝜇g/ml) (Fig. 1a). Both species tolerated the CTX treatment, which reduced the biomass production of *Klebsiella* and *Pseudomonas* by 46% and 26%, respectively, when measured after 27 h (Fig. 1a, bottom). *Klebsiella* grew ∼ 1.3-times faster than *Pseudomonas* without antibiotics (during the first 6 h) (Fig. 1a, top). With the CTX treatment, *Klebsiella* growth became detectable only after ∼ 11 h of incubation (Fig. 1a, bottom), while *Pseudomonas* growth was virtually unaffected (Fig. 1a).

**Figure 1:**
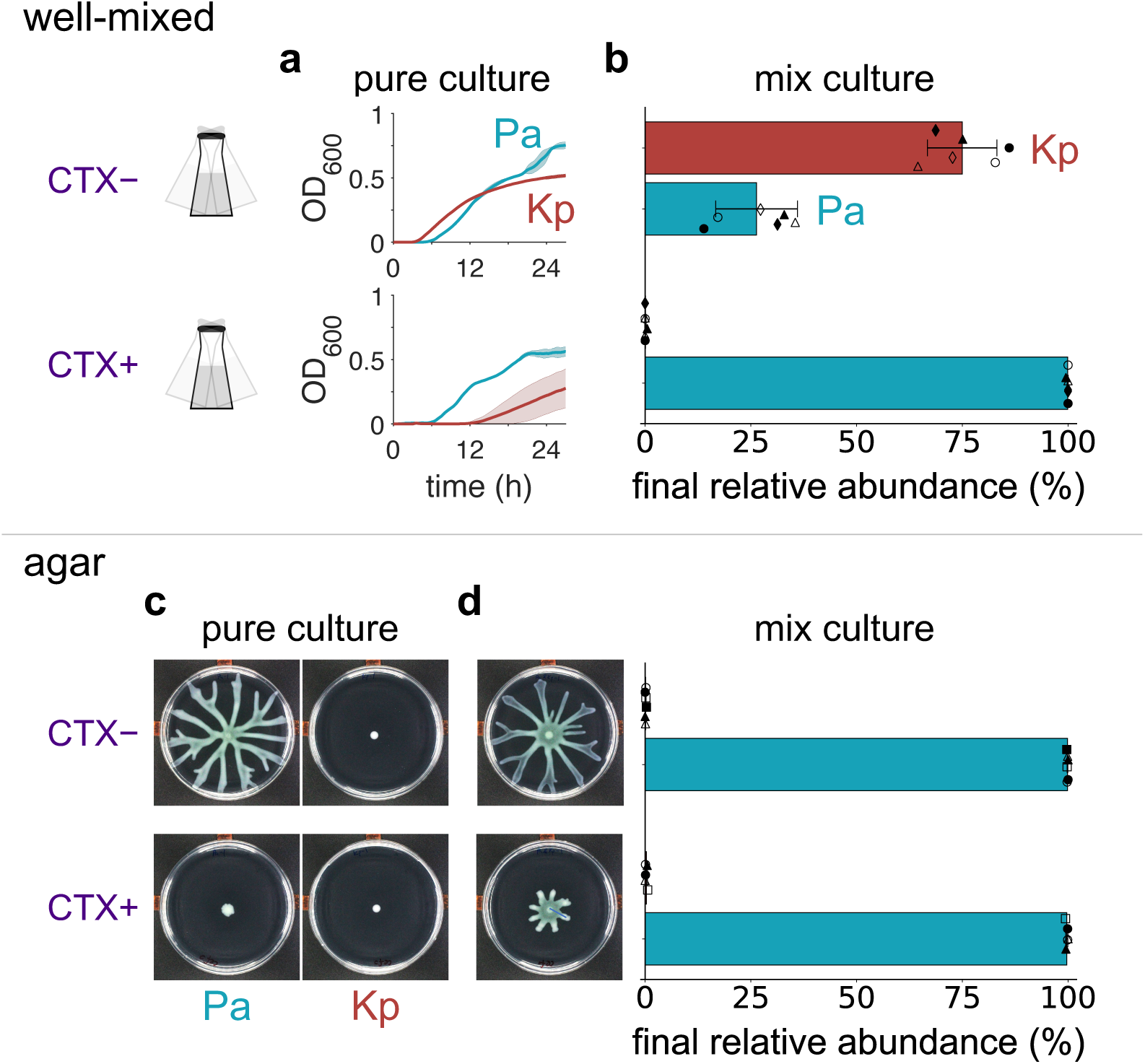
*Klebsiella* and *Pseudomonas* exhibited collective range expansion under CTX treatment at the cost of *Klebsiella*’s growth. *Klebsiella* and *Pseudomonas* pure cultures were grown overnight in lysogeny broth (LB). The cultures were then centrifuged, and the resulting cell pellets were resuspended in phosphate-buffered saline (PBS) supplemented with 8 g/l of casamino acids (see Methods). Kp: *Klebsiella*, Pa: *Pseudomonas*. **a. *Pseudomonas* tolerated CTX treatment better than *Klebsiella*.** *Well-mixed* pure cultures of *Klebsiella* (dark red) and *Pseudomonas* (cyan) were inoculated at an initial OD_600_ = ∼ 2.1×10_-5_ (estimated based on dilutions applied to a cell suspension with OD_600_ > 0.05) and without or with CTX (2.5 𝜇g/ml) (top and bottom, respectively). Curves and shades represent the average and the standard deviation, respectively, from two biological replicates each with three technical replicates. See Methods for details. **b. *Pseudomonas* dominated *Klebsiella* under the CTX treatment in well-mixed co-cultures.** Experiments were performed similarly to that in Fig. 1a. The liquid growth media was inoculated with a 1:1 volume mixture of the two species at an initial OD_600_ = ∼ 2.1×10_-5_. Co-cultures were incubated for 27 h and then harvested to measure relative abundances by plating (Supplementary Fig. 4b). Circles, triangles, and diamonds represent independent replicates whose technical replicates were shown by solid or open markers. Data represented by diamonds were obtained using fluorescently labeled strains (*Pseudomonas*-YFP and *Klebsiella*-mCherry). Data and error bars represent the mean and the standard deviation from the independent experiments after their technical replicates were averaged. When grown as a well-mixed pure culture, each species produced less biomass in medium conditioned by the other species than in unconditioned medium, demonstrating competition between the two species (Supplementary Fig. 2). **c. *Pseudomonas* range expansion was suppressed by CTX and rescued by *Klebsiella*.** 20 ml of swarming agar (0.55 %) media surface was center inoculated with 1 𝜇l of a 1:1 volume mixture of *Klebsiella* and *Pseudomonas* (OD_600_ = ∼ 0.43). Colony images were captured at 24 h, processed, analyzed for colony areas using a custom Python script (Methods; Supplementary Fig. 3). Standard 100-mm petri dishes were used. The images were equally adjusted for brightness and contrast for presentation. See Supplementary Fig. 7 for raw images from all replicates. **d. The rescue of the coculture range expansion occurred at the expense of *Klebsiella*’s growth.** After being imaged at 24 h, whole colonies were harvested by flooding with 5 ml of saline. The relative abundances of the two species in the harvests were estimated based on their distinct CFU morphologies (Supplementary Fig. 4b). Circles, triangles, and squares show independent experiments, for which the solid and open markers represent technical replicates. The data and error bars represent the mean and the standard deviation from the independent experiments after their technical replicates were averaged. Standard 100-mm petri dishes were used. The images were equally adjusted for brightness and contrast for presentation. See Supplementary Fig. 7 for raw images from all replicates. Also, see Supplementary Fig. 4a for the total biomasses of the cell harvests as well as its distribution between *Klebsiella* and *Pseudomonas*, estimated based on their relative abundances.

The results above show that *Klebsiella* grew faster without antibiotic treatment, whereas *Pseudomonas* grew faster under the CTX treatment. When grown as a well-mixed pure culture, each species produced less biomass in medium conditioned by the other species than in unconditioned medium, demonstrating competition between the two species (Supplementary Fig. 2). If the two species primarily interact through competition in well-mixed cocultures, we expect the dominance of *Klebsiella* without the CTX treatment and the dominance of *Pseudomonas* with the CTX treatment. To this end, we tested the growth of mix cultures, each starting with equal fractions of *Klebsiella* and *Pseudomonas*, with or without CTX treatment. Indeed, without CTX treatment, *Klebsiella* reached a relative abundance of 75.0% after 27 h (Fig. 1b, top). With CTX (2.5 𝜇g/ml) treatment, *Pseudomonas* reached a relative abundance of 99.9% after 27 h (Fig. 1b, bottom).

We then examined the two species and their interactions on an agar surface where *Pseudomonas* could exhibit swarming motility^52^. We tested the effect of the β-lactam antibiotic CTX (2.5 𝜇g/ml) on pure cultures of *Pseudomonas* or *Klebsiella* by inoculating them on agar and measuring their colonized areas and biomasses after 24 h of incubation (Fig. 1c; Supplementary Fig. 3, Supplementary Fig. 4a). The CTX treatment suppressed *Pseudomonas* growth by 28-fold when measured by colony areas (Fig. 1c; Supplementary Fig. 3), or 22-fold when measured by biomass (Supplementary Fig. 4a). This result is consistent with the studies that showed that *Pseudomonas*’ swarming is inhibited by antibiotic treatment^53,54^. In contrast, the CTX treatment did not significantly affect *Klebsiella* colony growth (Fig. 1c; Supplementary Fig. 3; Supplementary Fig. 4a). In both cases, *Klebsiella* colonies were smaller than those of *Pseudomonas* colonies (Fig. 1c; Supplementary Fig. 3).

We next measured the growth of a coculture starting with equal biomasses of the two species under the CTX treatment. The coculture colonized a 7.4-fold larger area and produced 6.2-fold greater biomass compared to *Pseudomonas* alone (Fig. 1c,d; Supplementary Fig. 3; Supplementary Fig. 4a). Without antibiotic treatment, *Pseudomonas* alone grew better than a mixture (Fig. 1c,d; Supplementary Fig. 3; Supplementary Fig. 4a), though the difference was statistically insignificant (A two-sample t-test assuming unequal variances (Welch’s t-test) was used to test the null hypothesis of no difference. The corresponding two-tailed *P*–values are 0.081 and 0.25 for colonized area and biomass, respectively). The collective range expansion by *Klebsiella* and *Pseudomonas* under the CTX treatment was also observed when different incubation temperatures (30 °C and 40 °C instead of the 37 °C default) were tested (Supplementary Fig. 5).

When mixed colonies were harvested and the relative abundances of the two species were measured based on their colony forming unit (CFU) morphologies (Supplementary Fig. 4b), we found that *Pseudomonas* dominated in cocultures with (99.7%) or without (99.9%) CTX treatment (Fig. 1d). Distributing the biomass in the mixed colonies with respect to these relative abundances, we found that *Klebsiella* always suffered from the mixed inoculation, while *Pseudomonas* benefited from it under the CTX treatment (Supplementary Fig. 4a, compare the colored bars with and without a grey frame).

To facilitate the analysis of the colony composition in a spatially resolved manner, we used fluorescence proteins to label the two species (*Pseudomonas*-YFP and *Klebsiella*-mCherry). In a coculture consisting of these two labeled strains, we observed that *Klebsiella* was mostly confined at the seeding position, whereas the expanded colony front largely consisted of *Pseudomonas* (Supplementary Fig. 6).

Collectively, our results indicate that the CTX treatment suppressed *Pseudomonas*’ spatial expansion, which was rescued by *Klebsiella* in a co-culture (Fig. 1c,d). In turn, the activated expansion of *Pseudomonas* suppressed the growth of *Klebsiella* (Fig. 1d; Supplementary Fig. 4a), which could be explained by a greater tolerance of *Pseudomonas* to CTX than *Klebsiella* (Fig. 1a).

The rescuing by *Klebsiella* was likely due to its ability to produce Bla, which degrades CTX^45^ (Supplementary Table 1). To test this notion, we collected supernatants from the *Klebsiella* pure cultures at ∼ 6.3 h or media without cell inoculation as unconditioned control. After filter sterilizing the supernatants, we tested whether CTX-sensitive *Escherichia coli* could grow in them (Supplementary Fig. 8a, top). Following 21 h of incubation, *E.* coli exhibited detectable growth in the *Klebsiella*-supernatant but not in the unconditioned control, consistent with the CTX degradation by *Klebsiella*-produced Bla^45^ (Supplementary Fig. 8a, bottom; see Supplementary Fig. 8b, top for *Klebsiella* growth curves with the initial cell density of OD_600_ = ∼ 5×10^-4^, which was used for this experiment).

### CTX suppressed *Pseudomonas*’ spatial expansion in a dose-dependent manner

We next measured the suppression of *Pseudomonas* spatial expansion (in terms of colonized area) using varied initial doses of CTX (0 to 2.5 𝜇g/ml) (Supplementary Fig. 9). The suppression increased with the initial CTX dose (*A*_0_), approximately following a Hill inhibition function.

*Pseudomonas* growth was virtually unaffected by CTX (2.5 𝜇g/ml) treatment (Fig. 1a). Thus, the suppression in *Pseudomonas*’ spatial expansion (Fig. 1c; Supplementary Fig. 9) was likely due to the loss of motility and correlated with filamentous cell morphology (Supplementary Fig. 10), consistent with previous reports associating β-lactam-induced cell filamentation with reduced motility^55,56^. The extent of the filamentation of *Pseudomonas* cells was reduced when inoculated in a mixture with *Klebsiella* (Supplementary Fig. 10), which restored *Pseudomonas*’ spatial expansion (Fig. 1d). A similar filamentation of *Pseudomonas* was observed in well-mixed liquid cultures under CTX (2.5 𝜇g/ml) treatment, with or without *Klebsiella* (Supplementary Fig. 11). The spatial expansion of *Pseudomonas* was eventually stopped by the antibiotic concentration far (1.5 ∼ 2 cm) away from the inoculation point. This is because *Klebsiella* did not move along with *Pseudomonas* (Supplementary Fig. 6) and, hence, could no longer reduce the antibiotics to an ineffective concentration. Likewise, actively swarming *Pseudomonas* still needed *Klebsiella* for swarming when inoculated on fresh antibiotic media at ∼ 4-times greater cell density than the original inoculum (Supplementary Fig. 12).

### Theory reveals key determinants of the keystone engineering effect

We conducted theoretical analysis to gain insight into the *Klebsiella*-mediated collective range expansion during antibiotic treatment. We assumed that the *Klebsiella* population leads to an antibiotic ‘sink’ and the growth media serves as a ‘source’ for antibiotics (Fig. 2a, top)^57^. The terms ‘sink’ and ‘source’ were inspired by the ‘infinite source & point sink’ reaction-diffusion framework we adapted below. We described the dependence of the effective spatial expansion ability (*M*) of *Pseudomonas* on the antibiotic concentration using the experimentally determined Hill function (Supplementary Fig. 9). Solving the system assuming cylindrical symmetry (see Supplementary Information for details), we have:

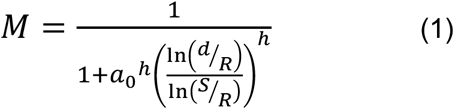

where 𝑅 > 0 is the boundary of the ‘sink’ created by the *Klebsiella* population, 𝑆 ≥ 𝑅 is the domain boundary, 𝑑 ∈ [𝑅, 𝑆] defines the distance of *Pseudomonas* from the ‘sink’ boundary, and 𝑎_0_ ∈ [0, ∞] is the initial antibiotic concentration normalized by its value for the half-maximum of *Pseudomonas*’ effective spatial expansion ability (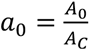; see Supplementary Fig. 9 caption for 𝐴_𝐶_ and ℎ). Here, the ‘sink’ boundary *R* defines the radius of a circular ‘clear zone’ within which the antibiotic concentration is an ineffective lower boundary value. For the latter, we considered zero concentration for simplicity, though our theoretical framework can be applied to any arbitrary value. That is, each point on the ‘clear zone’ boundary acts as a ‘point sink’ for the antibiotics.

**Figure 2:**
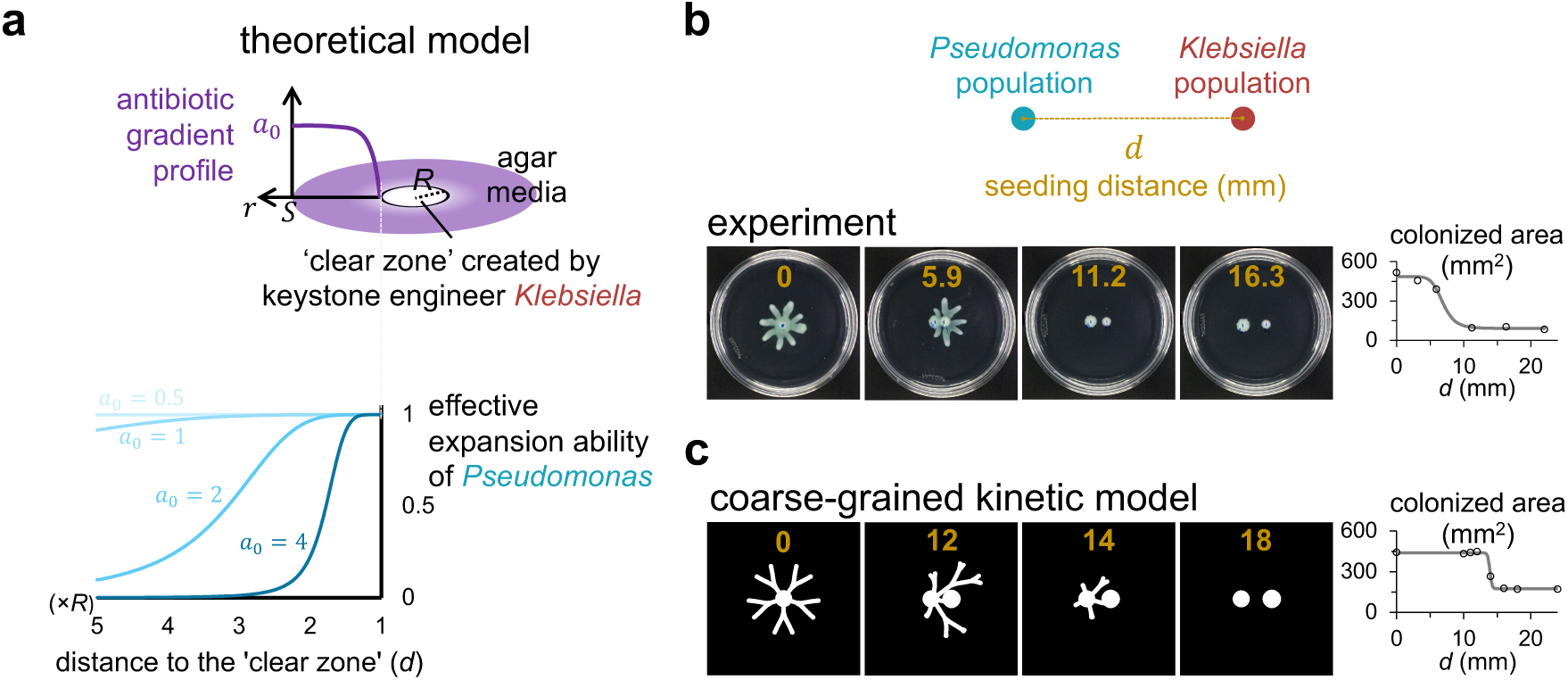
Spatial scale of the keystone engineering role. **a. Theoretical model: *Top*: ‘*Klebsiella*–antibiotic’ interaction**. Antibiotic concentration (𝑎) has a steady-state gradient, under the assumption that a population of the keystone engineer *Klebsiella* acts as an antibiotic sink creating a ‘clear zone’ and the growth media serves as an infinite antibiotic source. 𝑎 is smaller when closer to the ‘sink’, where the antibiotic concentration is minimized at an ineffective value (zero for simplicity; see Supplementary Information for details). ***Bottom*: ‘*Klebsiella*-antibiotic-*Pseudomonas*’ interactions.** Equation 1 predicts that *Pseudomonas*’ effective spatial expansion ability (*M*) is determined by its distance to the sink boundary (*d*) and the initial antibiotic concentration (𝑎_0_). *h* = 6.642 (Supplementary Fig. 9), and *S*/*R* = 10 was assumed. **b. Experiments corroborated the dependence of *Pseudomonas*’ effective spatial expansion ability on its distance to *Klebsiella.*** The experiment in Fig. 1d with CTX (2.5 𝜇g/ml) was repeated with the only difference being 1 𝜇l from the *Pseudomonas* (left) and 1 𝜇l from the *Klebsiella* (right) cell suspensions were inoculated side-by-side with various distance between them (gold in mm). The dishes were imaged at 23 h. The colonized areas were measured using a custom Python script and manually corrected using ImageJ when needed. The data was fitted to a Hill inhibition function (solid curve), where the Hill coefficient and the spatial scale of the keystone engineering (𝐿_𝐾_) are 7.62 and 6.90 mm, respectively (𝑅^2^= 0.990). Each population initially occupied a circular area with a radius of ∼ 1.5 mm. Standard 100-mm petri dishes were used. The images were equally adjusted for brightness and contrast for presentation. See Supplementary Fig. 13 for the raw data from all replicates and Supplementary Fig. 14 for the data from an experiment with the fluorescently labeled strains. Note that the half-maximum suppressing CTX dose was 1.29 μg/ml (Supplementary Fig. 9). **c. A coarse-grained kinetic model recapitulated the experimental range expansion patterns.** The results from the coarse-grained simulation at 48 h. *Left*. Black and white indicates bacteria are absent and present, respectively. *Right*. Total area where bacteria present was calculated using a custom MATLAB script and plotted as colonized area as a function of the seeding distance (center-to-center) between the *Pseudomonas* and *Klebsiella* populations (gold in mm). The data was again approximated by a Hill inhibition function (solid curve), where the Hill coefficient and 𝐿_𝐾_ are 92.4 and 13.9 mm, respectively (𝑅^2^= 0.999). Each population initially occupied a circular area with a radius of 5 mm. See Supplementary Fig. 15 for the data at different time points. See Supplementary Table 2 for model parameters and initial conditions.

Note that we expect the ‘clear zone’ radius, *R*, to have two contributions: it scales with the physical size of the *Klebsiella* inoculum and has an additional term that increases nearly linearly with the characteristic diffusion length scale of Bla. This additional contribution is anticipated to depend only weakly on the *Klebsiella* seeding cell density (see Supplementary Information, Equation S7 for details).

For a constant 𝑑, 𝑀 decreases from 1 and approaches 0 with increasing 𝑎_0_. For a given 𝑎_0_, 𝑀 decreases with increasing *d* (Fig. 2a, bottom). For a sufficiently large 𝑎_0_, *Pseudomonas*’ effective spatial expansion ability is sharply activated when *d* < *L_K_*, where *L_K_* denotes the distance at which 𝑀 is half-maximum. Thus, *L_K_* is a measure of the spatial scale of keystone engineering. Critically, *L_K_* can be substantially larger than *R*, indicating a long-range interaction (Fig. 2a, bottom, *L_K_* = ∼ 3.2*R* for 𝑎_0_ = 2).

### Experiments corroborated the theoretical predictions

To test Equation 1, we inoculated one *Klebsiella* population and one *Pseudomonas* population at varied distance (𝑑) under CTX (2.5 𝜇g/ml) treatment. Then, we quantified the range expansion as the colonized areas at 23 h, observing a sharp change at a certain distance value (Fig. 2b), consistent with the prediction of the spatial scale of the keystone engineering (𝐿_𝐾_) above. The range expansion had a bias towards the *Klebsiella* population, which we confirmed by repeating this experiment with the fluorescently labeled strains (Supplementary Fig. 14). We demonstrated the robustness of the existence of 𝐿_𝐾_, repeating the same experiment, also with varied *Klebsiella* seeding cell density (Supplementary Fig. 13).

### A coarse-grained kinetic model recapitulated the experimental range expansion patterns

The spatial patterns of the range expansion (Fig. 2b) were not considered in our theoretical estimation of 𝑀 and 𝐿_𝐾_ above. The spatial patterns are a function of not only the antibiotic gradient and the *Pseudomonas*’ spatial expansion ability—which the antibiotic gradient determines—but also the spatiotemporal dynamics of species and nutrient abundances. Thus, we next accounted for these factors and built a coarse-grained model, enabling a more constrained simulation of spatial expansion dynamics (Methods and Supplementary Table 2).

This model builds upon our previous models for *Pseudomonas* branching^58^ and Bla-mediated population dynamics in response to β-lactam treatment^45^ (see Methods). We simulated the range-expansion dynamics with varied seeding distance (𝑑) between one *Klebsiella* population and one *Pseudomonas* population. Again, we quantified the total colonized area (at 48 h; see Supplementary Fig. 15 for different time points) and found a sharp onset of range expansion as 𝑑 decreased below a critical value, consistent with the existence of 𝐿_𝐾_ (Fig. 2c). Simulations mimicking an intracellular degradation of antibiotics by an immotile species (zero dispersal for the antibiotic-degrading enzyme and the species that produces it) yielded similar results, suggesting that the ‘antibiotic sink’ role could be fulfilled without released public goods (Supplementary Fig. 16).

### The keystone engineering occurred in a pairwise community of hospital sink co-isolates, *Bacillus paranthracis* and *Pseudomonas aeruginosa*

Colonization of hospital sink drains (e.g., p-traps) has been associated with pathogens such as *K. pneumoniae*^59^ and *P. aeruginosa*^60^. Moreover, biofilms formed in the p-traps of connected sinks and expanded upwards 2.5 cm/day, possibly due to surface-associated bacterial motility^61^. If sufficiently close to the sink strainer, running sink water can lead to the spread of bacteria to the environment^61^, posing a health risk.

We next tested whether the keystone engineering could occur in communities of bacteria isolated from the same sink p-trap at Duke University Hospital. Two isolates, 192621 and 192622, were identified by 16S rDNA sequencing as *Bacillus paranthracis* and *P. aeruginosa*, respectively.

We first tested the effect of carbenicillin (CB; 10 𝜇g/ml) treatment on the growth of the two isolates. Specifically, we grew *B. paranthracis* 192621 (Bp) and *P. aeruginosa* 192622 (Pa22) in pure liquid cultures without or with CB (10 𝜇g/ml) (Fig. 3a). Both species tolerated the CB treatment, but Pa22 grew faster than Bp under both conditions. When grown as a well-mixed pure culture, each species produced less biomass in medium conditioned by the other species than in unconditioned medium, demonstrating competition between the two species (Supplementary Fig. 17). If the two species primarily interact through competition in well-mixed cocultures, we expect the dominance of Pa22 under both conditions. To this end, we tested the growth of mix cultures, each starting with equal fractions of Bp and Pa22, with or without CB treatment. Indeed, Pa22 dominated after 27 h for both cases, where Bp was undetectable (< 0.1%) (Fig. 3b).

**Figure 3:**
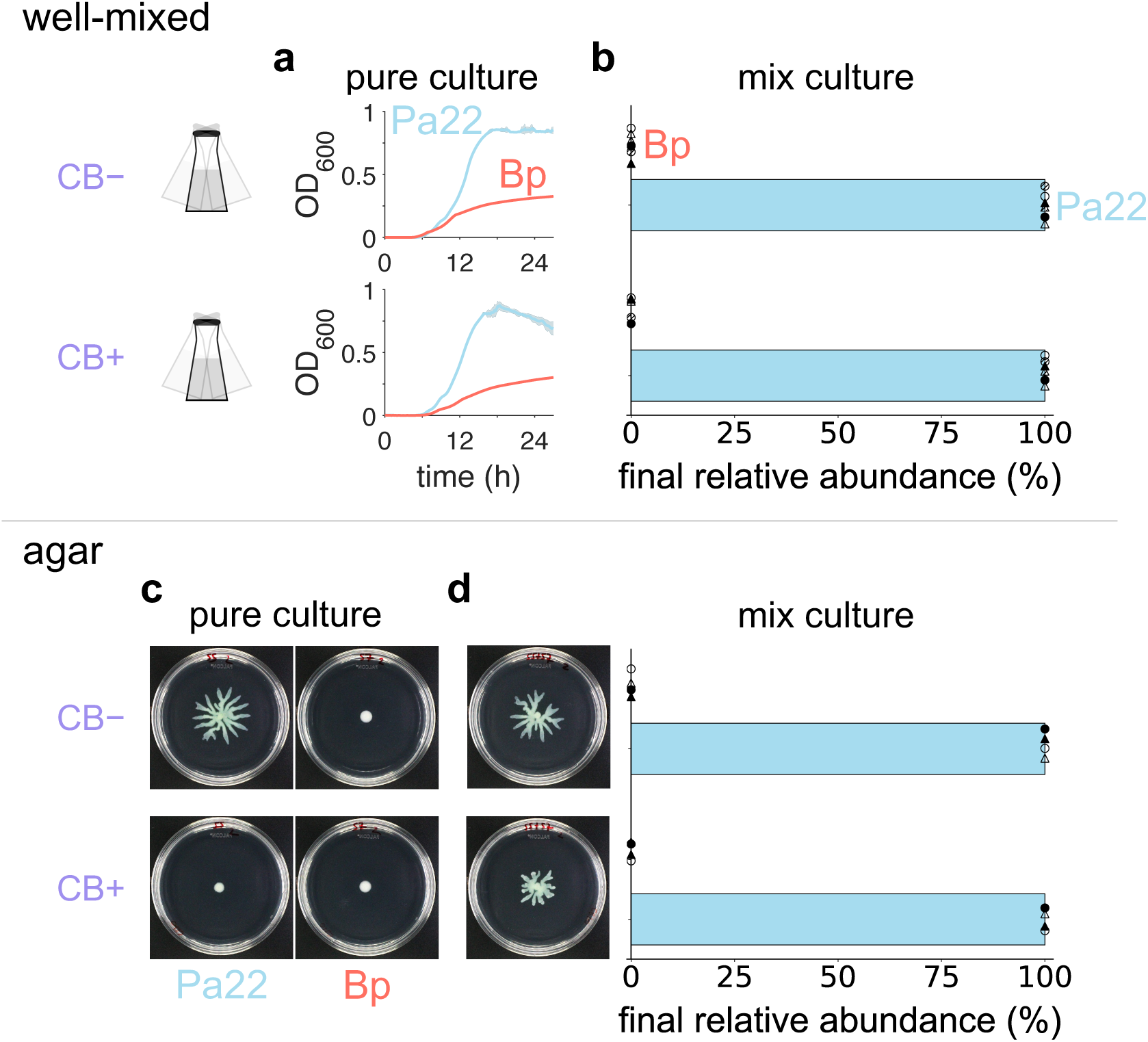
The keystone engineering in a pairwise community of hospital sink isolates, Bp and Pa22. Bp and Pa22 pure cultures were grown overnight in LB. The cultures were then centrifuged, and the resulting cell pellets were resuspended in PBS supplemented with 8 g/l of casamino acids (see Methods). Bp: *Bacillus paranthracis*, Pa22: *Pseudomonas aeruginosa* 192622. **a. Pa22 tolerated CB treatment better than Bp.** *Well-mixed* cultures of Bp (red) and Pa22 (light blue) were inoculated, with the initial cell density of OD_600_ = ∼ 2.1×10_-5_ (estimated based on dilutions applied to a cell suspension with OD_600_ > 0.05), and without or with CB (10 𝜇g/ml) (top and bottom, respectively). Curves and shades represent the average and the standard deviation, respectively, from two biological replicates each with three technical replicates. **b. Pa22 dominated Bp in well-mixed cocultures via competition.** The liquid growth media was inoculated with a 1:1 volume mixture of the two species at the starting cell density of OD_600_ = ∼ 2.1×10_-5_. Co-cultures were incubated for 27 h and then harvested to measure relative abundances by plating (Supplementary Fig. 20b). Circles and triangles represent independent replicates whose technical replicates were shown by solid, open, or lined markers. Data and error bars represent the mean and the standard deviation from the independent experiments after their technical replicates were averaged. When grown as a well-mixed pure culture, each species produced less biomass in medium conditioned by the other species than in unconditioned medium, demonstrating competition between the two species (Supplementary Fig. 17). **c. Pa22’s range expansion was suppressed by CB treatment and rescued by Bp.** 20 ml of agar (0.55 %) media surface was center inoculated with 1 𝜇l of a 1:1 volume mixture of Bp and Pa22 with or without CB (10 μg/ml) at the initial cell density of OD_600_ = ∼ 0.34 or ∼ 0.43). Images of the colonies were captured at 24 h. The images were processed and analyzed for colonized areas using a custom Python script (Methods). Standard 100-mm petri dishes were used. The images were equally adjusted for brightness and contrast for presentation. See Supplementary Fig. 23 or all raw images. **d. The rescue of the coculture range expansion occurred at the expense of Bp’s growth.** After being imaged at 24 h, whole colonies were harvested by flooding with 5 ml of saline. The total biomass was calculated by multiplying the absorbance (OD_600_) and the volume of the suspension. The relative abundances of the two species in the harvests were estimated based on their distinct CFU morphologies (Supplementary Fig. 20b). Circles (inoculum density of OD_600_ = ∼ 0.43) and triangles (inoculum density of OD_600_ = ∼ 0.34) represent independent experiments, for which the solid and open markers show technical replicates. Data and error bars show the mean and standard deviation, respectively, from all the replicates. Standard 100-mm petri dishes were used. The images were equally adjusted for brightness and contrast for presentation. See Supplementary Fig. 23 or all raw images. Also, see Supplementary Fig. 20a for the total biomasses of the cell harvests as well as its distribution between Bp and Pa22, estimated based on their relative abundances.

When tested on agar, however, only a mixture of Bp and Pa22 exhibited spatial expansion after 24 h under treatment by CB (10 μg/ml; ∼ 2.4-times the concentration for the half-maximum suppression of the spatial expansion of Pa22 when alone, Supplementary Fig. 18) (Fig. 3c,d). The mixture’s spatial expansion under the CB treatment was maintained at 30 °C but was not at 40 °C (Supplementary Fig. 19), indicating the context-dependence of mechanism. While mixing Pa22 with Bp promoted biomass production under the CB treatment by 4-fold, Bp itself became undetectable (< 0.1%) in the final population (Fig. 3d; see Supplementary Fig. 20 for biomass data). Supernatants from CB-treated pure cultures of Bp allowed the growth of CB-sensitive *E. coli* (Supplementary Fig. 21), indicating the degradation of CB in the environment by Bp. These results suggest that Bp served the keystone engineer role on Pa22 only during community spatial expansion by degrading the antibiotics in the environment.

The principles we uncovered from the pairwise communities above (Fig. 1,2,3) are likely generalizable: When we used a variant of the CB-sensitive *E. coli*, which was engineered to express Bla conferring CB resistance^62,63^, it also rescued *Pseudomonas* swarming from CB inhibition (Supplementary Fig. 22).

### Keystone engineering occurred in a synthetic community of sink isolates

Pairwise communities above enabled us to investigate mechanistic underpinnings of the keystone engineering that enables collective range expansion under antibiotic treatment. To evaluate the generalizability of the mechanism, we next tested if Bp’s keystone engineering on Pa22’s spatial expansion could occur in the presence of other members of sink isolates. To this end, we assembled an eight-member community, which we termed ‘SynkC’, by introducing six additional isolates from the same sink sample from which we isolated both Bp and Pa22. SynkC consists of six distinct taxonomic groups based on 16S rDNA-based microbiome sequencing. Seven out of the eight SynkC members are genotypically and/or phenotypically distinct (Supplementary Fig. 24). In addition to Pa22, only one other SynkC member showed spatial expansion ability on its own (Supplementary Fig. 24a, *Bacillus sp.* ESY92). When tested alone, two SynkC members did not tolerate the CB treatment, and the others exhibited various degrees of tolerance (Supplementary Fig. 24b).

Whole genome sequencing of the SynkC members revealed that each member had at least one Bla gene. Bp had the most (17) Bla genes followed by Pa22 (7), while Bp was the only isolate carrying a class A Bla, according to Ambler’s classification scheme, based on amino acid sequence similarities of known Bla proteins (Supplementary Table 1). These sequencing results suggest a greater or a unique potential of Bp to degrade certain β-lactam antibiotics. Accordingly, in pairwise cultures, Bp was the only one that rescued the Pa22 spatial expansion from the CB treatment inhibition (Supplementary Fig. 25).

We ran agar experiments with SynkC with equal starting biomasses of all the eight members or by leaving out Bp (SynkC-Bp). Again, spatial expansion under the CB treatment occurred only when Bp was included (Fig. 4, images), though Bp was not detected in the final population of SynkC (Fig. 4, pie charts). In addition to promoting the spatial expansion and ∼ 2.7-fold greater total biomass, the initial presence of Bp also increased community diversity under the CB treatment. Specifically, while only Pa22 was detected at the end of the SynkC-Bp experiment, both Pa22 and *Enterobacter sp*. were detected in the final SynkC community (Fig. 4, pie charts). This suggests that *Enterobacter sp.* should also be considered when analyzing the precise spatial distribution of bacteria. However, *Enterobacter sp.* cannot spread alone (Supplementary Fig. 24a), so any spatial of *Enterobacter sp*. must depend on other species such as Pa22.

**Figure 4:**
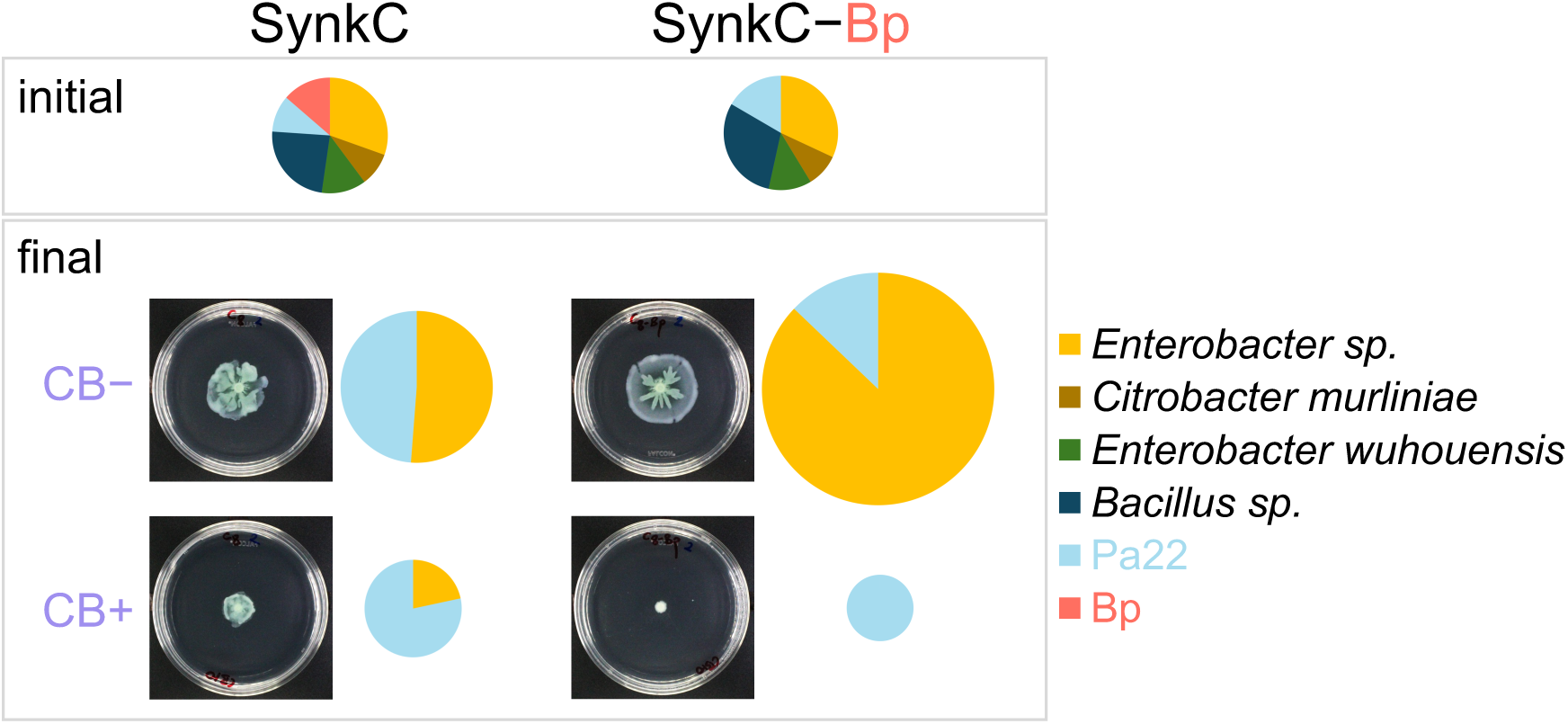
The keystone engineering in SynkC, an 8-member, synthetic community of hospital sink co-isolates.

The agar (0.55%) surface was inoculated with 1 𝜇l of a mixture containing equal biomasses of all the eight isolates (SynkC) or all but Bp (SynkC–Bp) (OD_600_ = ∼ 0.34). Images of the colonies were captured at 24 h. Then, whole colonies were harvested by flooding with ∼ 5 ml of saline. The biomasses of the cell harvests were calculated by measuring the absorbance (OD_600_) of each harvest suspension and multiplying it by the suspension volume. The initial and final cell suspensions were stored as frozen glycerol stocks and underwent 16S microbiome sequencing to determine the initial and final relative abundances of different community members (a 0.5 % cut-off value applied). Each of the pie charts above shows six groups, with the *Enterobacter sp.* and *Bacillus sp.* groups representing the combined abundance of two isolates each, while all other groups represent a single isolate. For the final time point, relative areas of the pie charts represent the distribution of the respective biomasses measured. Two independent experiments, each with technical replicates, yielded similar results. A representative set of data was shown here, after the images were equally processed for brightness and contrast for presentation. Standard 100-mm petri dishes were used. CB (10 𝜇g/ml) was used when applicable. See Supplementary Fig. 26 for the raw data from all replicates. See Supplementary Fig. 24 for the identities and phenotypes of the individual isolates.

## Discussion

Our study provides a concrete demonstration that context—specifically spatial structure and chemical stress—can reveal a species’ keystone engineer role. By degrading β-lactam antibiotics, an immotile Bla-producing species (*K. pneumoniae* or *Bacillus paranthracis*) creates a ‘clear zone’ that restores the motility-based range expansion of a more resilient partner (*P. aeruginosa*). The keystone engineering we report here represents a form of facilitation. *K. pneumoniae* and *B. paranthracis* actively modify the environment by degrading antibiotics, thereby enabling the range expansion of *P. aeruginosa*. However, this facilitation is at the cost of the facilitators themselves—driving them to an extreme minority population, apparently through competitive exclusion by *P. aeruginosa*. Although the keystone engineer eventually becomes a minority due to competition, its transient facilitation is essential for spatial expansion and overall community success. This mechanism could contribute to the observed rapid biofilm formation in sink-associated microbial communities^61^.

Our results broaden the canonical interpretation of keystone species, which typically emphasizes effects on growth or community structure^14–17,24,28–33^. Two contextual factors are critical for our system. One is spatial expansion, as mobility only matters in a spatially structured environment. This context-dependency is reminiscent of a recent example in which we show that the spatial expansion of *Pseudomonas* not only makes a cooperative trait essential^42^ but also allows non-cooperating variants (so-called “cheaters”) to leverage their growth advantage from the lack of cooperative trait expression, enabling them to outcompete the cooperating individuals^43^. The other contextual factor is antibiotic stress, which primarily suppresses mobility instead of the growth of *Pseudomonas*. Together, these factors shift the net effect of the keystone engineer from competitive to facilitative. This shift aligns with the stress gradient hypothesis, which suggests that positive interactions dominate negative interactions in more stressful environments^64–67^.

The principles uncovered here are likely generalizable. Other species or consortia could similarly modify their environment—through pH shifts^24^, enzyme secretion^68,69^, nutrient release/break-down^31,68,69^, or other chemical changes—to unlock traits like motility^70,71^, biofilm growth^72–74^, or nutrient utilization (i.e., necrotrophy^68^) relevant only in spatially structured habitats. Such facilitation may be common in natural and engineered microbial communities but remain untapped due to a lack of spatially explicit experimental inquiry.

Clinically, consistent with the notion of private versus public benefits of Bla producers in bacterial communities^75^, our work suggests that certain Bla producers may play a crucial, even if transient, role in promoting the spread of more virulent pathogens, such as *Pseudomonas*, even if they remain undetected in standard assays. Current microbiome analyses often exclude low-abundance taxa^76^, potentially missing these keystone engineers and underestimating their influence on infection dynamics.

In conclusion, recognizing the context-dependent emergence of keystone engineering enriches our understanding of microbial ecology, offers insights for controlling infections, and opens new avenues for studying hidden facilitators in complex microbial ecosystems.

## Methods

### Bacterial growth media and procedures

In every experiment, bacteria were first streaked on a lysogeny broth (LB) agar plate from their frozen glycerol stock, and a single colony was incubated overnight in liquid LB media at 37 °C and 225 r.p.m.

For all experiments, PBS (2.4 g/l Na_2_HPO_4_ (anhydrous), 3 g/l KH_2_PO_4_ (anhydrous), 0.5 g/l NaCl, 1 mM MgSO_4_, and 0.1 mM CaCl_2_) supplemented with casamino acids (8 g/l, unless otherwise noted) was used. Agar (final concentration of 0.55 %) was used when applicable.

The media was prepared following a recipe adapted from Xavier *et al*.^77^: To make 1 liter of 5x phosphate buffer stock solution, 12 g Na_2_HPO_4_ (anhydrous), 15 g KH_2_PO_4_ (anhydrous), and 2.5 g NaCl were dissolved in deionized water and sterilized by autoclaving. The casamino acids stock solutions were made at 20 % (w/v) by microwaving 40 g casamino acids (Gibco™ Bacto™ 223120) in deionized water, sterilized by filtering (0.22 μm) and stored at 4 °C. The main solutions of agar were made at 1.25 % (w/v) on the day of the experiment dissolving granulated agar (BD Difco™ 214530) in deionized water and sterilizing by autoclaving. Then, appropriate amounts were immediately used to prepare the experimental media.

In the experiments and overnight cultures with the *Pseudomonas*-YFP and *Klebsiella*-mCherry variants, gentamycin (15 μg/ml) was added for the maintenance of the plasmids. aTc (100 ng/ml) was also added for inducing the mCherry expression. All antibiotic solutions used in the study were made fresh on the day of the experiment.

All OD_600_ measurements were taken using 200 μl samples and a plate reader (Tecan Infinite 200).

#### Well-mixed liquid cultures

We grew 2-3 ml cells in 16 ml culture tubes at 37 °C and 225 r.p.m. The cultures with aTc were protected from light.

##### Growth curve experiments

We measured the growth curves of bacteria using a plate reader (Tecan Infinite 200). In each well of a 96-well plate, we added 195 μl liquid media and 5 μl cell inoculum culture (prepared by centrifuging overnight-grown cells at 1150 g for 5 min and then resuspending in the liquid medium). To prevent evaporation, we used Nunc Edge multi-well plates (Thermo Scientific) with built-in water reservoirs. In addition to the built-in water reservoirs, we also reserved all the edge wells for water only. The cells were then incubated in the plate reader at 37 °C, and OD_600_ measurements were taken at 10 min intervals after the plate being shaken rigorously. For technical replicates of a particular condition, cells from the same inoculum were inoculated in different wells of a 96-well plate and measured. Background signals measured from media containing no cells were subtracted from the data. Custom MATLAB scripts were used for data analysis, during which a four time point moving average was applied to the raw OD_600_ time series data.

#### Agar media experiments

20 ml growth media was solidified in 100 mm petri dishes. Overnight-grown cells were centrifuged at 1150 g for 5 min and then resuspended in the liquid medium to prepare a cell inoculum culture. The seeding droplet volume from cell inoculum cultures was 1 μl. The seeded plates were incubated at 37 °C, agar side up.

#### CTX/CB environmental degradation assays

The protocol used was adapted with minimal changes from our previous study^45^: Briefly, 3 ml liquid media with/without the keystone engineer bacteria (*Klebsiella* or Bp) inoculated (initial cell density of OD_600_ = ∼ 5×10^-4^) and with/without the antibiotic (2.5 𝜇g/ml CTX or 10 𝜇g/ml CB) addition were incubated at 37 °C and 225 r.p.m. At ∼ 6.3 h, 1 ml samples from the media were removed, in which clavulanic acid (5 μg/ml final concentration) was then added to prevent any further Bla activity. Next, supernatants were prepared by filtration of these 1 ml samples using 0.22 𝜇m cellulose acetate filters (VWR). Finally, the supernatants were inoculated with *E. coli* MC4100Z1 at initial cell density of OD_600_ = ∼ 5×10^-4^, at which the *E. coli* is sensitive to the antibiotic treatments here. Then, OD_600_ measurements were taken after 21 h incubation at 37 °C at 225 r.p.m.

#### Spent media assays

Similar to the “CTX/CB environmental degradation assays” above, 3 ml liquid culture media were incubated for ∼ 6.3 h at 37 °C and 225 r.p.m. with no inoculant (for an unconditioned control media) or with *Klebsiella* or *Pseudomonas* (or with Bp or Pa22 at the initial cell densities odf OD_600_ = ∼ 5×10^-4^). Then, supernatants were collected by filtration using 0.22 μm cellulose acetate filters (VWR). Within each pairwise system, the filtered supernatants of each species were cross-inoculated with the other species (OD_600_ = ∼ 5×10^-4^). Subsequently, growth curve measurements were performed in a plate reader at 37 °C, as described above.

#### Relative abundance measurements based on colony forming unit (CFU) morphology

As shown in Supplementary Fig. 4b and Supplementary Fig. 20b, CFUs were grown on LB agar (1.5 %) for 20 h at 37 °C to determine the relative abundance of the constituent species in pairwise mixed cultures. To do so, cell harvests were serially diluted in sterile saline (11.6 g/l NaCl) solution down to ∼ 100 CFUs per 100 μl spread on a 100-mm LB agar plate.

### Bacterial strains

*P. aeruginosa* PA14 (*Pseudomonas*), a clinical *K. pneumoniae* isolate D-005 from Deverick J. Anderson’s laboratory (*Klebsiella*), *E. coli* MC4100Z1, *E. coli* MC4100Z1 with a plasmid that encodes an engineered cytoplasmic β-lactamase (*E. coli* Bla+)^62^, and the hospital sink p-trap isolates described below were used. For some experiments variant strains that expressed spectrally distinct fluorescent proteins from a plasmid, *Pseudomonas*-YFP^43^ and *Klebsiella*-mCherry (this study, see Methods and Supplementary Table 3), were used.

#### Construction of the Klebsiella-mCherry strain

##### Strain & transformation

Zymo Mix & Go Transformation kit (Zymo Research) was used to make *K. pneumoniae* D-005 chemically competent with one modification: the overnight culture of D-005 was re-grown in super optimal broth (SOB) supplemented by 0.7 mM (final concentration) ethylenediaminetetraacetic acid (EDTA)^78^.

##### Plasmid construction

ptetmCherryGMR was constructed from the parent plasmids ptetmCherry^79^, pUCP30T-eCFP plasmid^80^, and pJ1996_v2^81^. The ptetmCherry plasmid without the chloramphenicol resistance gene, the gentamicin resistance gene, and a DNA fragment containing two transcription terminators were PCR amplified from the ptetmCherry, pUCP30T-eCFP, and pJ1996_v2 plasmids, respectively, by DNA primers with overhangs. The resulting linear DNA fragments were digested by DpnI and subsequently column purified with DAN Clean & Concentor-5 (Zymo Research). Then, they were ligated together with NEBuilder HiFi DNA Assembly Master Mix (NEB #E2621) to yield the final plasmid. The final plasmid was transformed into Top10F’ to propagate, miniprepared with Zymo Plasmid Miniprep-Classic kit (Zymo Research) and sequenced by Sanger sequencing (Genewiz/Azenta Life Sciences) to validate.

All the DNA primers were purchased from Integrated DNA Technologies and listed in Supplementary Table 3.

#### SynkC: Isolation of bacteria from the sink p-traps and their taxonomic identification

| Strain identity | Taxonomic group name we used | Identity suggested by the Azenta Life Sciences pipeline | Identity suggested by the SeqCenter LLC pipeline |
| --- | --- | --- | --- |
| ESY55 | Pa22 | *Pseudomonas aeruginosa*<br>ATCC10145 | *Pseudomonas aeruginosa* |
| ESY57 | Bp | *Bacillus paranthracis*<br>MCCC1A00395 | *Bacillus* |
| ESY66 | *Enterobacter wuhouensis* | *Enterobacter wuhouensis/ludwigii* | *Enterobacter cloacae* |
| ESY73 | *Bacillus sp.* | *Bacillus aerophilus/stratosphericus/altitudinis* | *Bacillus sp.* |
| ESY80 | *Citrobacter murlinae* | *Citrobacter murlinae / freundii* | *Citrobacter* |
| ESY83 | *Enterobacter sp.* | *Enterobacter bugandensis* 247BMC | *Enterobacter* |
| ESY92 | *Bacillus sp.* | *Bacillus aerophilus/stratosphericus/altitudinis* | *Bacillus sp.* |
| ESY93 | *Enterobacter sp.* | *Enterobacter sichuanensis/asburiae* | *Enterobacter* |

The members of SynkC were isolated by propagations of a sample collected from a sink p-trap in an intensive care unit in a recently constructed bedtower at the Duke University Hospital. Sterile tubing was fed down into the p-trap and a 50 ml syringe was used to agitate the fluid to collect biofilm portions and then withdraw a sample^82^. 50 μl of p-trap water was used to inoculate 950 μl of tryptic soy broth in a 2 ml deep well plate (Thermo Scientific) and grown for 16 h at 37 °C with shaking at 700 r.p.m. The resulting culture was glycerol stocked (25% glycerol) and frozen. In parallel, it was serially diluted to 10^-6^ in phosphate buffered saline and 10 μl of dilutions from 10^-3^ to 10^-6^ were plated on tryptic soy agar (TSA) with and without 100 μg/ml carbenicillin. The plates were incubated at 37 °C for 10 h and at room temperature for 66 h.

Pa22 (ESY55) and Bp (ESY57) were isolated via restreaking different parts of the resulting colonies on fresh agar plates and incubating for 18 h at 37 °C. A new set of overnight cultures were then made by inoculating 2 ml of tryptic soy broth with individual colonies on the restreaked plates and incubating for 16 hours at 37 °C. These cultures were then glycerol stocked as the eventual Pa22 and Bp. Subsequent colonies from these stocks were sent for 16S sequencing at Azenta Life Sciences for species identification.

ESY66 and ESY73 were isolated as follows: First, the glycerol stock of the original overnight culture described above was inoculated LB and then grown for ∼ 8 h at 37 °C with shaking at 225 r.p.m. The resulting culture was glycerol stocked (25% glycerol) and frozen. This stock is called the “master stock” hereafter. The master stock was diluted ∼ 10^4^-fold in 11.6 g/l saline, and 100 μl of the dilution was plated on TSA plates. The plates were incubated at 37 °C for ∼ 20 h. Individual colonies were restreaked on LB agar plates, and the plates were incubated at 37 °C for ∼ 20 h, again. Then, overnight cultures were grown by inoculating 2 ml of LB with individual colonies from the restreaked plates and incubating for ∼ 18 h at 37 °C with shaking at 225 r.p.m. These cultures were then glycerol stocked as the eventual isolates used in this study. Individual colonies from these stocks were sent for 16S sequencing at Azenta Life Sciences for species identification.

ESY80, ESY83, ESY92, and ESY93 were isolated as follows: First, ∼ 1 μl from the master stock was inoculated on swarming agar media and incubated for ∼ 24 h at 37 °C. The final colony was suspended in 11.6 g/l saline and then glycerol (25 %) stocked and frozen. This new frozen stock was diluted ∼ 10^5^-fold in 11.6 g/l saline, and 100 μl of the dilution was plated on TSA plates. The plates were incubated at 37 °C for ∼ 20 h. Individual colonies were restreaked on LB agar plates, and the plates were incubated at 37 °C for ∼ 20 h, again. Then, overnight cultures were grown by inoculating 2 ml of LB with individual colonies from the restreaked plates and incubating for ∼ 18 h at 37 °C with shaking at 225 r.p.m. These cultures were then glycerol stocked as the eventual isolates used in this study. Individual colonies from these stocks were sent for 16S sequencing at Azenta Life Sciences for species identification.

#### 16S microbiome sequencing by SeqCenter

16S V3/V4 region sequencing (20000 reads) service of SeqCenter, LLC was used. Aliquots from cell harvests were supplemented by glycerol (25 % final concentration) before being frozen at -80 °C. The frozen aliquots were shipped overnight to the company’s facility on dry ice.

##### DNA extraction

All standard DNA extractions at SeqCenter follow the ZymoBIOMICS™ DNA Miniprep Kit. 20 - 200 μl from the samples submitted were transferred into 550 μl of lysis solution. Cells suspended in lysis solution were transferred into the ZR BashingBead™ Lysis Tubes and mechanically lysed using the MP FastPrep-24™ lysis system with 1 min of lysis at maximum speed and 5 min of rest for 5 cycles. Samples were then centrifuged at 10000 g for 1 min. 400 μl of supernatant was transferred from the ZR BashingBead™ Lysis Tube to a Zymo-Spin™ III-F Filter and centrifuged at 8000 g for 1 min. 1200 μl of ZymoBIOMICS™ DNA Binding Buffer was added to the effluent and mixed via pipetting. 800 μl of this solution was transferred to a Zymo-Spin™ IICR Column and centrifuged at 10000 g for 1 min. This step was repeated until all material was loaded onto the Zymo-Spin™ IICR Column. DNA bound to the Zymo-Spin™ IICR Column was washed 3 times with 400 μl and 700 μl of ZymoBIOMICS™ DNA Wash Buffer 1 and then 200 μl of ZymoBIOMICS™ DNA Wash Buffer 2 with a 1-min spin down at 10000 g for each, respectively. Washed DNA was eluted using 75 μl of ZymoBIOMICS™ DNase/RNase Free Water following a 5-min incubation at room temperature and a 1-min spin down at 10000 g. The Zymo-Spin™ III-HRC Filter was prepared using 600 μl of the ZymoBIOMICS™ HRC Prep Solution and a centrifugation at 8000 g for 3 min. Eluted DNA was then purified by running the effluent through the prepared Zymo-Spin™ III-HRC Filter. Final DNA concentrations were determined via Qubit.

##### Sequencing

Samples were prepared using Zymo Research’s Quick-16S kit with phased primers targeting the V3/V4 regions of the 16S gene. The specific primer sequences are found below:

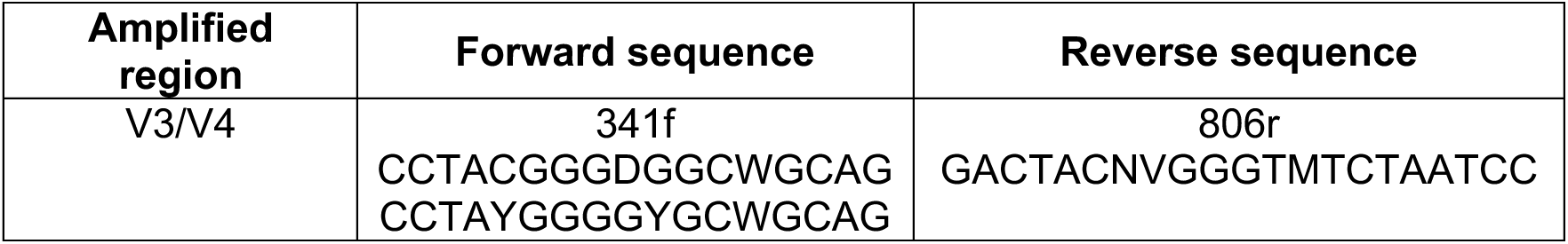

Following clean up and normalization, samples were sequenced on a P1 600cyc NextSeq2000 Flowcell to generate 2×301 bp paired end (PE) reads. Quality control and adapter trimming was performed with bcl-convert (v4.2.4).

##### Analysis

Sequences were imported to Qiime2^83^ for analysis. Primer sequences were removed using Qiime2’s cutadapt^84^ plugin using the following degenerate primer queries:

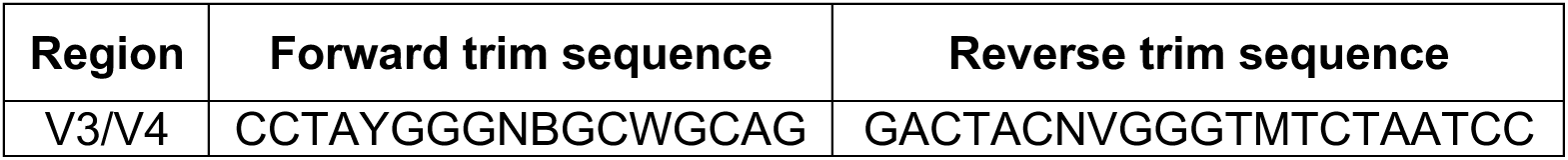

Sequences were then denoised using Qiime2’s dada2 plugin^85^. Denoised sequences were assigned operational taxonomic units (OTUs) using the “Silva 138 99% OTUs full-length sequence database” and the VSEARCH^86^ utility within Qiime2’s feature-classifier plugin. OTUs were then collapsed to their lowest taxonomic units, and their counts were converted to reflect their relative frequency within a sample.

Note that, for SynkC, neither the SeqCenter microbiome sequencing nor the Azenta Life Sciences 16S sequencing-based species identification pipelines could distinguish the members within the *Enterobacter sp*. and *Bacillus sp* taxonomic groups. The identities of Pa22, Bp, *Citrobacter murliniae*, and *Enterobacter wuhouensis* were better resolved to species level from the best fits of the Azenta Life Sciences pipeline, so we used those identities. We confirmed that these four isolates did not fall into either the *Enterobacter sp*. or the *Bacillus sp.* taxonomic groups in the SeqCenter pipeline, which was used to generate the relative abundance data in Fig. 4 and Supplementary Fig. 26.

#### Whole genome sequencing by SeqCenter LLC

##### DNA extraction

DNA extractions were performed by SeqCenter LLC following the ZymoBIOMICS™ DNA Miniprep Kit: A loopful of cells (∼ 50 – 100 mg) were aseptically scraped from the submitted agar plates and resuspended in 750 μl of lysis solution.

Cells suspended in lysis solution were transferred into the ZR BashingBead™ Lysis Tubes and mechanically lysed using the MP FastPrep-24™ lysis system with 1 min of lysis at maximum speed and 3 min of rest for 2 cycles. Samples were then centrifuged at 10,000 rcf for 1 min. 400 μl of supernatant was transferred from the ZR BashingBead™ Lysis Tube to a Zymo-Spin™ III-F Filter and centrifuged at 8,000 rcf for 1 min. 1,200 μl of ZymoBIOMICS™ DNA Binding Buffer was added to the effluent and mixed via pipetting. 800 μl of this solution was transferred to a Zymo-Spin™ IICR Column and centrifuged at 10,000 rcf for 1 min. This step was repeated until all material was loaded onto the Zymo-Spin™ IICR Column.

DNA bound to the Zymo-Spin™ IICR Column was washed 3 times with 400 μl and 700 μl of ZymoBIOMICS™ DNA Wash Buffer 1 and then 200 μl of ZymoBIOMICS™ DNA Wash Buffer 2 with a 1-min spin down at 10,000 rcf for each, respectively. Washed DNA was eluted using 75 μl of ZymoBIOMICS™ DNase/RNase Free Water following a 5-min incubation at room temperature and a 1-min spin down at 10,000 rcf. The Zymo-Spin™ III-HRC Filter was prepared using 600 μl of the ZymoBIOMICS™ HRC Prep Solution and a centrifugation at 8,000 rcf for 3 min. Eluted DNA was then purified by running the effluent through the prepared Zymo-Spin™ III-HRC Filter. Final DNA concentrations were determined via Qubit.

##### Illumina sequencing

Illumina sequencing libraries were prepared using the tagmentation-based and PCR-based Illumina DNA Prep kit and custom Integrated DNA Technologies 10 bp unique dual indices with a target insert size of 280 bp. No additional DNA fragmentation or size selection steps were performed. Illumina sequencing was performed on an Illumina NovaSeq X Plus sequencer in one or more multiplexed shared-flow-cell runs, producing 2×151 bp paired-end reads. Demultiplexing, quality control and adapter trimming was performed with bcl-convert (a proprietary Illumina software for the conversion of bcl files to basecalls, v4.2.4).

##### Illumina short-read assembly and annotation

Short read assembly was performed with Unicycler^87^. Assembly statistics were recorded with QUAST^88^. Samples were annotated with Bakta^89^.

| Tool | Version | Parameters |
| --- | --- | --- |
| unicycler | 0.5.0 | default parameters |
| quast | 5.2.0 | default parameters |
| bakta | 1.8.1 | default parameters; db version 5.0 |

### Imaging

Bacterial colonies on agar plates were imaged with a UVP Colony Doc-It Imaging Station with epi white light, using the same brightness and contrast settings for all images. The same z-plane focus was applied to all images within the same independent experiment.

Microscopic imaging was performed using a Keyence BZ-X710 microscope and Keyence BZX Software Suite 1.3.1.1. To image a whole swarming colony, its entire plate was scanned and imaged, and the resulting images were stitched using the built-in algorithm of the software.

### Image analyses

A custom Python script was used for processing and analyzing the colonized area for bacterial colony images. This script firsts finds and removes the plate boundaries. Then, the bacterial colony (or colonies when applicable) is automatically segmented using a user-defined threshold intensity value. Finally, the image background is changed to plain white for display purposes. When applicable, the script calculated the seeding distance between different species as the Euclidean distance between two positions, defined by the user based on the distinct appearance of the inoculum zones. For each case, the segmentation results were manually validated and corrected using Image J manual tools when necessary.

Custom MATLAB scripts were used for the coarse-grained kinetic model simulations and the corresponding analyses.

### Coarse-grained kinetic model

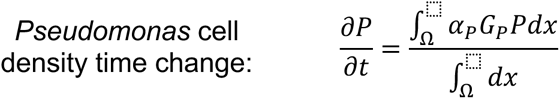

where 𝑃 is *Pseudomonas* cell density (cell density unit: cu), 𝛼_𝑃_ is the growth rate constant of *Pseudomonas* (1/h), Ω indicates the entire colony domain, 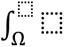 indicates an integral over the entire colony domain. 𝐺_𝑃_ is the *Pseudomonas* growth function as defined below,

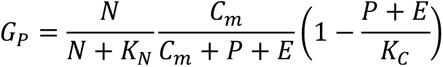

where 𝑁 is the local nutrient (casamino acids) concentration (g/l), 𝐾_𝑁_ is the local nutrient concentration for half-maximum growth rate (g/l), 𝐶_𝑚_ is the local cell density for the half-maximum suppression of the growth rate (cu), 𝑃 is the local *Pseudomonas* cell density (cu), 𝐸 is the local *Klebsiella* cell density (cu), and 𝐾_𝐶_ is the local carrying capacity (cu).

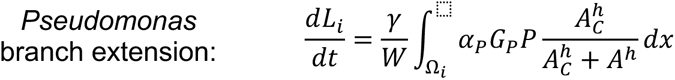

where 𝐿_𝑖_ is the length of branch 𝑖 (mm), 𝛾 *Pseudomonas* spatial expansion efficiency (mm/(h.cu)), 𝑊 is the branch width previously measured from experimental data (mm)^58^, 𝐴 is the local cefotaxime antibiotic concentration (mol/l), 𝐴_𝐶_ is cefotaxime concentration for half-maximum suppression of *Pseudomonas* effective spatial expansion ability (mol/l), ℎ is the steepness of the suppression of *Pseudomonas* effective spatial expansion ability by cefotaxime, Ω_𝑖_ indicates the domain of branch 𝑖, and 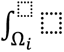 indicates an integral over branch 𝑖 domain.

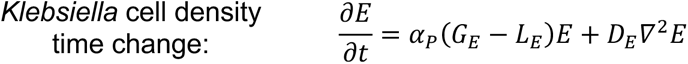

where 𝐷_𝐸_ is the diffusive dispersal rate of *Klebsiella* (mm^2^/h), and 𝐺_𝐸_ and 𝐿_𝐸_ are *Klebsiella* growth and lysis functions, respectively, as defined below.

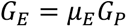

where 𝜇_𝐸_ is the growth rate constant of *Klebsiella* in proportion to that of *Pseudomonas*.

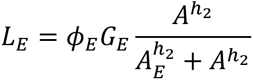

where 𝜙_𝐸_ is the proportionality constant between the growth and lysis rates of *Klebsiella*, 𝐴_𝐸_ is cefotaxime concentration for half-maximum *Klebsiella* lysis rate (mol/l), and ℎ_2_ is the steepness of the onset of *Klebsiella* lysis by cefotaxime.

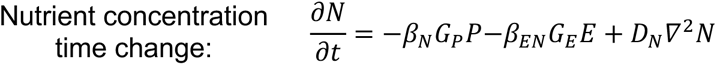

where 𝛽_𝑁_ and 𝛽_𝐸𝑁_ are the rate constants for nutrient consumption by *Pseudomonas* and *Klebsiella*, respectively, cell density growth (g/(l.h.cu) for both). Note that 𝛽_𝐸𝑁_ must be constrained tightly as *Pseudomonas* would otherwise avoid *Klebsiella*-colonized territory due to nutrient depletion. 𝐷_𝑁_ is the diffusivity of nutrients (mm^2^/h).

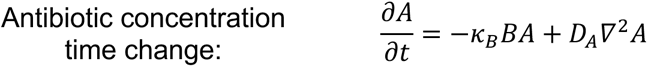

where 𝐵 is the local concentration of extracellular Bla (mol/l), 𝜅_𝐵_ is the rate constant for cefotaxime antibiotic degradation by Bla (l/(mol.h)), and 𝐷_𝐴_ is the diffusivity of antibiotics (mm^2^/h).

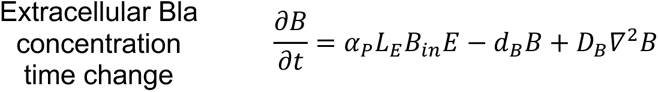

where 𝐵_𝑖𝑛_ is the intracellular concentration of Bla (mol/cell; for *Klebsiella* only), 𝑑_𝐵_ is the spontaneous degradation rate constant for Bla (1/h), and 𝐷_𝐵_ is the diffusivity of Bla (mm^2^/h).

We have formulated the above mathematical model building upon our previous models for *Pseudomonas* branching pattern formation^58^ and Bla-producing bacteria population dynamics under β-lactam treatment^45^:

As we lack a complete mechanistic understanding of how *Pseudomonas* colonies develop branches, our model defines the colony morphology formation based on a set of explicit rules, as described by Luo, *et al*.^58^: (1) A colony initiates from a circular inoculum. (2) Branches emerge with a defined branch width *W* and branch density *D* (number of branch tips per unit area), both measured from experimental data. (3) Each branch 𝑖 extends at an extension rate 𝑑𝐿_𝑖_⁄𝑑𝑡, which is proportional to cell growth within the branch. (4) Each branch extends following the local nutrient gradient (towards where the nutrient concentration increases most steeply) and bifurcates when the local cell density reaches the threshold *D*.

At each timestep, we compute cell growth, nutrient consumption, antibiotics degradation, etc., and determine the extension rate from cell growth using the equations described above. For simplicity, we assume that the cell density redistributes and becomes uniform within the colony at each timestep.

Based on this framework, we incorporated antibiotic inhibition into the branch extension to account for the effect of antibiotics on colony expansion. Also, from our previous model for Bla-producing bacteria population dynamics under β-lactam treatment^45^, we incorporated *Klebsiella* growth and lysis due to antibiotics, Bla production by lysing *Klebsiella*, and antibiotic degradation by Bla.

We used the parameter values from the two original papers^45,58^, with some minor modifications, when possible. The diffusivities of the antibiotic and Bla were chosen based on their molecular weights relative to nutrients (i.e., casamino acids), as suggested by Einstein’s relation. The parameter values for the cefotaxime concentration for the half-maximum suppression of *Pseudomonas*’ effective spatial expansion ability (𝐴_𝐶_) and the sharpness of this suppression (ℎ) were used from the Hill-fits to our experimental data above (see Supplementary Fig. 9a caption).

We assumed no spontaneous degradation of the antibiotic or nutrient recycling after lysis. Finally, we added another parameter to account for our experimental observation above that *Pseudomonas* spatial expansion is effectively biased towards *Klebsiella* (Fig. 2b and Supplementary Fig. 14). This parameter essentially determines the relative dominance of *Pseudomonas*’ range expansion on nutrient gradients versus antibiotic gradients depending on their global profiles: *Pseudomonas* would move towards decreasing antibiotic concentration (i.e., the *Klebsiella* neighborhood), except when 𝑑 = 0, if the spatial global average of antibiotic concentration is greater than 𝐴_𝐶_ and if the global minimum of nutrient concentration is greater than a factor (𝐴_<_ = 0.7 as the default value) of the nutrient concentration for half maximum specific growth rate (𝐾_𝑁_). For 𝑑 = 0, this factor was applied as 𝐴_<_ = 10^16^ to represent an infinitely large value.

The partial differential equations were numerically solved using a custom MATLAB implementation of the alternating direction implicit method by applying no-flux boundary conditions. The numerical simulations were run in a 1001-by-1001-pixel spatial domain, where each pixel dimension was equivalent to a ∼ 0.106 mm spatial length based on the model parameters used. Each population was started in a circular zone equivalent of 5 mm radius. The simulation timestep used was equivalent to 0.02 h, again, based on the model parameters used.

See Supplementary Table 2 for the values of the model parameters as well as the initial conditions used.

## Acknowledgments

We thank Lawrence A. David, Joanna B. Goldberg, David K. Karig, Jeffrey Letourneau, Jia Lu, Alice S. Prince, and David S. Weiss for helpful feedback. We thank Yasa Baig, Minsu Kim, Philip N. Rather, and Stuart A. West for their critical reading of an earlier version of the manuscript. This work was partially supported by grants from National Institutes of Health (LY: R01GM098642), Office of Naval Research (LY: N00014-20-1-2121), Defense Advanced Research Projects Agency (LY: 2501-203-2015861), National Science Foundation (LY & CTL: MCB-1937259), and by the Engineering Research Centers Program of the National Science Foundation under NSF Cooperative Agreement No. EEC-2133504. The funders had no role in study design, data collection and analysis, decision to publish, or preparation of the manuscript.

## Author contributions

E. Ş.: Conceived the study, designed, and conducted the research. Wrote the manuscript. C. A. V.: Assisted with experiments in Fig. 1 and Supplementary Fig. 10. K. S.: Assisted with experiments in Supplementary Fig. 13. Z. Z. Assisted with experiments in Supplementary Fig. 8a and Supplementary Fig. 21a. Wrote the cylindrically symmetric antibiotic gradient solution in the Supplementary Information. N. L.: Made observations that inspired this study. D. L.: Constructed the fluorescent protein-labeled variant of *Klebsiella* (*Klebsiella*-mCherry). Assisted in the experiments that led to Supplementary Fig. 10. H. R. M.: Assisted in the isolation of the hospital sink bacteria. D. J. A.: Obtained and provided the p-trap water samples which led to the isolation of the hospital sink bacteria. C. T. L.: Assisted with data interpretation, writing the manuscript, and articulating the broader ecological context. L. Y.: Conceived the study and assisted with research design and data interpretation. Wrote the manuscript.

All authors contributed to the reviewing and editing of manuscript drafts and approved the final version.

## Conflicts of interest

None are declared.

## Data availability

Sequencing data supporting Fig. 4, Supplementary Fig. 26, and Supplementary Table 1 are available at https://10.5281/zenodo.15882541. Data supporting Supplementary Fig. 5,9,10,13,14,19,24,25 are also available at https://10.5281/zenodo.15882541. All other data are provided in the main text, supplementary material, or at https://github.com/youlab/keystone.

## Code availability

Codes are available at https://github.com/youlab/keystone.

## Notes

### Competing Interest Statement

The authors have declared no competing interest.

### Summary of Updates

Text updated to clarify terminology. Main figures updated for enhanced clarity. New Supplementary Figures and Tables were added to address reviewers' comments. Corresponding authors information updated.

