## Supplementary Information for "Keystone engineering enables collective range expansion in microbial communities"

### Supplementary Figures

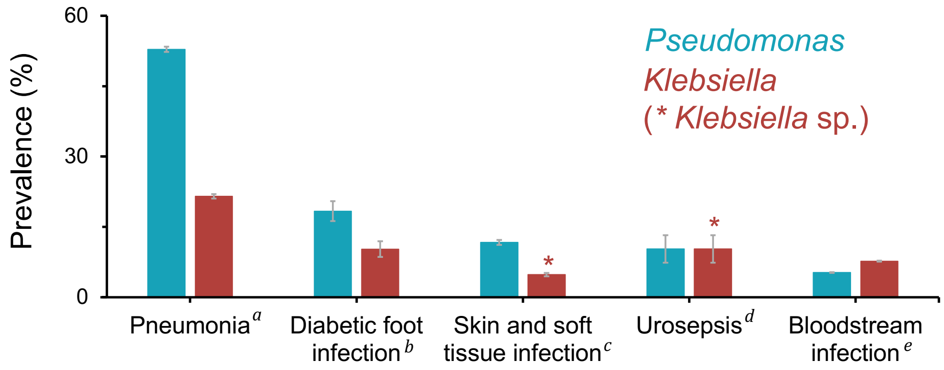


**Supplementary Figure 1: *Klebsiella* and *Pseudomonas* are prevalent in various infection types.**

We searched the literature to determine the prevalence *Klebsiella* (*K. pneumoniae*) and *Pseudomonas* (*P. aeruginosa*) in various infections. We selected data with sample numbers available only: pneumonia ($n$ = 28 918), diabetic foot infection ($n$ = 1 284), skin and soft tissue infection ($n$ = 12 920), urosepsis ($n$ = 408), and bloodstream infection ($n$ = 264 901). *Pseudomonas* was consistently among the top four prevalent bacterial species. *Klebsiella* was also prevalent in all categories, with skin and soft tissue infections and urosepsis data exceptionally including all *Klebsiella* species (*Klebsiella* sp.; indicated by '*'), which ranked as the sixth and second most prevalent bacterial entity, respectively. Note that urinary tract infections are often assessed with a corresponding blood infection. Hence, urosepsis represents cases where the same infecting pathogen was found in both urinary tract and blood samples. The error bars represent 95% confidence intervals calculated using the formula $1.96\sqrt{p(1-p)/n}$, where $n$ is the number of samples and $p$ is the fraction of samples identified as *Pseudomonas* or *Klebsiella*.

[a] Data from the SENTRY Antimicrobial Surveillance Program. One bacterial isolate per patient was collected from patients hospitalized with pneumonia in Western Europe, Eastern Europe, and the USA between 2016 – 2019[1].

[b] Data from a review that reported combined results of 10 studies from 2004 – 2018. Polymicrobial infections were represented in the data, but neither their overall percentage nor their species-level composition was reported[2].

[c] Data from the SENTRY Antimicrobial Surveillance Program. Isolates were collected from patients hospitalized with skin and soft tissue infections in North America, Latin America, and Europe between 1998 – 2004. No information was available for the number of isolates collected per patient[3].

[d] Global (70 countries included from four continents) patient data from 2003 – 2013. No information was found for the number of isolates collected per patient[4].

[e] Data from the SENTRY Antimicrobial Surveillance Program. One bacterial isolate per patient was collected from patients hospitalized with pneumonia in North America, Latin America, Europe, and the Asia-Pacific region between 1997 – 2016[5].

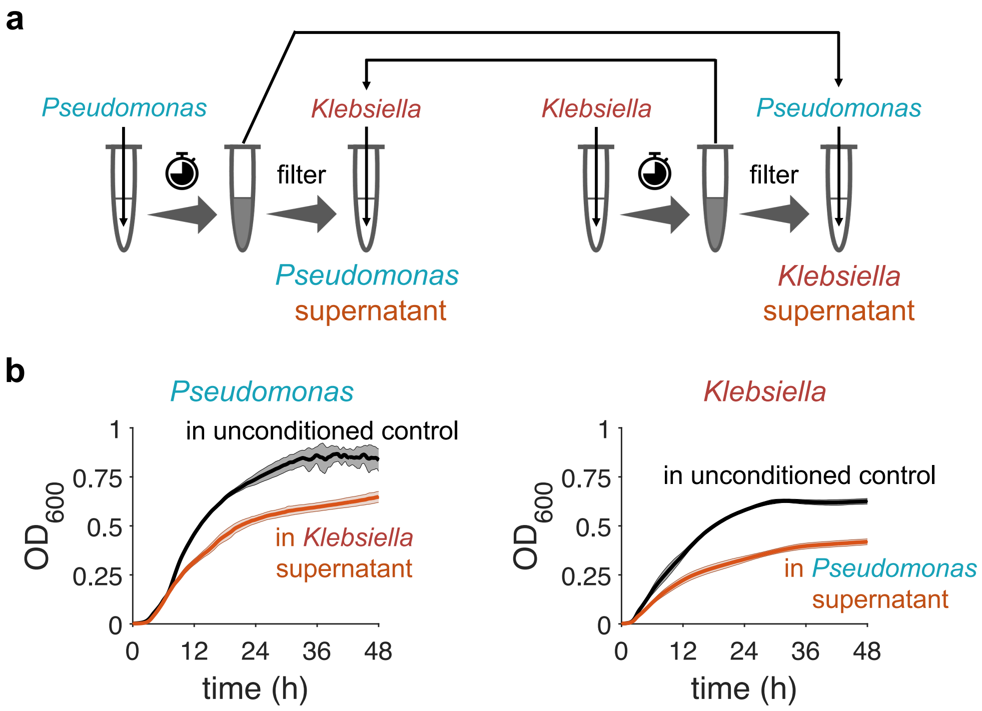


**Supplementary Figure 2: Spent media experiments demonstrated that *Pseudomonas* and *Klebsiella* primarily compete.**

**a.** Culture media were incubated for ~ 6.3 h at 37 °C and 225 r.p.m. with no inoculant (for an unconditioned control media) or with *Klebsiella* or *Pseudomonas* (initial cell density of $OD_{600}$ = ~ $5\times10^{-4}$). Then, supernatants were collected by filtration using 0.22 $\mu$m cellulose acetate filters (VWR). The filtered supernatants of each species were cross-inoculated with the other species ($OD_{600}$ = ~ $5\times10^{-4}$).

**b.** $OD_{600}$ measurements were taken in a plate reader at 37 °C. For each case, 12 replicates were performed—two independent replicates each with two technical replicates for the prior growth phase and three technical replicates for the spent media growth phase. The curves and shades represent the average and the standard deviation, respectively, from all replicates.

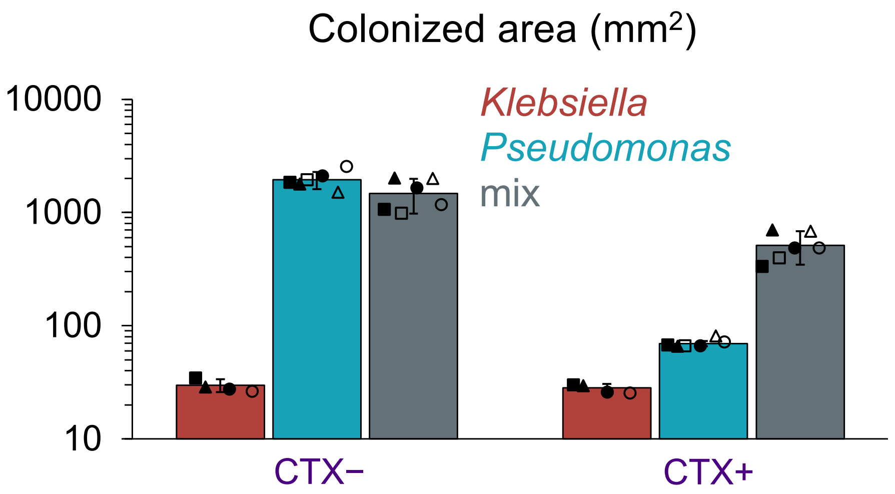


**Supplementary Figure 3: The quantification of the colonized areas in Fig. 1c,d.**

The images in Fig. 1c,d and Supplementary Fig. 7 were analyzed for colonized areas using a custom Python script (Methods). See Supplementary Fig. 7 for raw images.

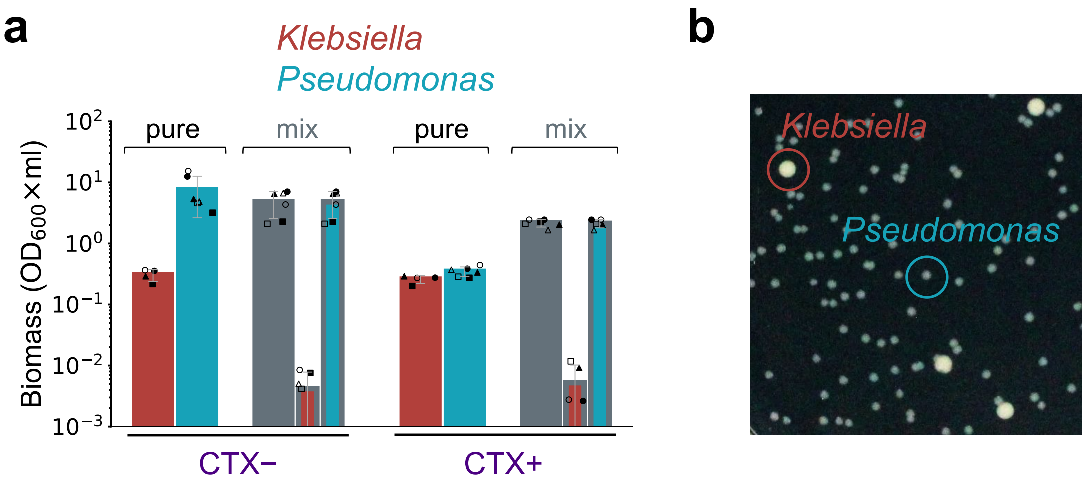


**Supplementary Figure 4: *Klebsiella* and *Pseudomonas* biomass production in pure- and mix- cultures on agar media, in which the latter was distributed based on the distinct colony forming unit (CFU) morphologies of the two species.**

**a.** The total biomass of the cell harvests in Fig. 1c,d and Supplementary Fig. 7 was calculated by measuring the absorbance ($OD_{600}$) of the harvest suspension and multiplying it by the harvest's volume. Relative contributions of *Klebsiella* and *Pseudomonas* to the total biomass values for "mix" (the colored bars with a grey frame) were estimated by multiplying the total "mix" biomass with the relative abundances of *Klebsiella* and *Pseudomonas* (Fig. 1d), determined by plating based on distinct CFU morphologies of the two species. Circles, triangles, and squares represent biological replicates whose technical replicates were shown by solid or open markers. The data and error bars represent the mean and standard deviation, respectively, from the biological replicates after their technical replicates were averaged.

**b. Distinct CFU morphologies of *Klebsiella* and *Pseudomonas*.** We distinguished between *Klebsiella* and *Pseudomonas* by plating them on LB agar (1.5 %) plates and categorizing them the following day based on their distinct CFU appearances. Here we show an example image, processed for brightness and contrast, where *Klebsiella* (dark red) and *Pseudomonas* (cyan) colonies are outlined.

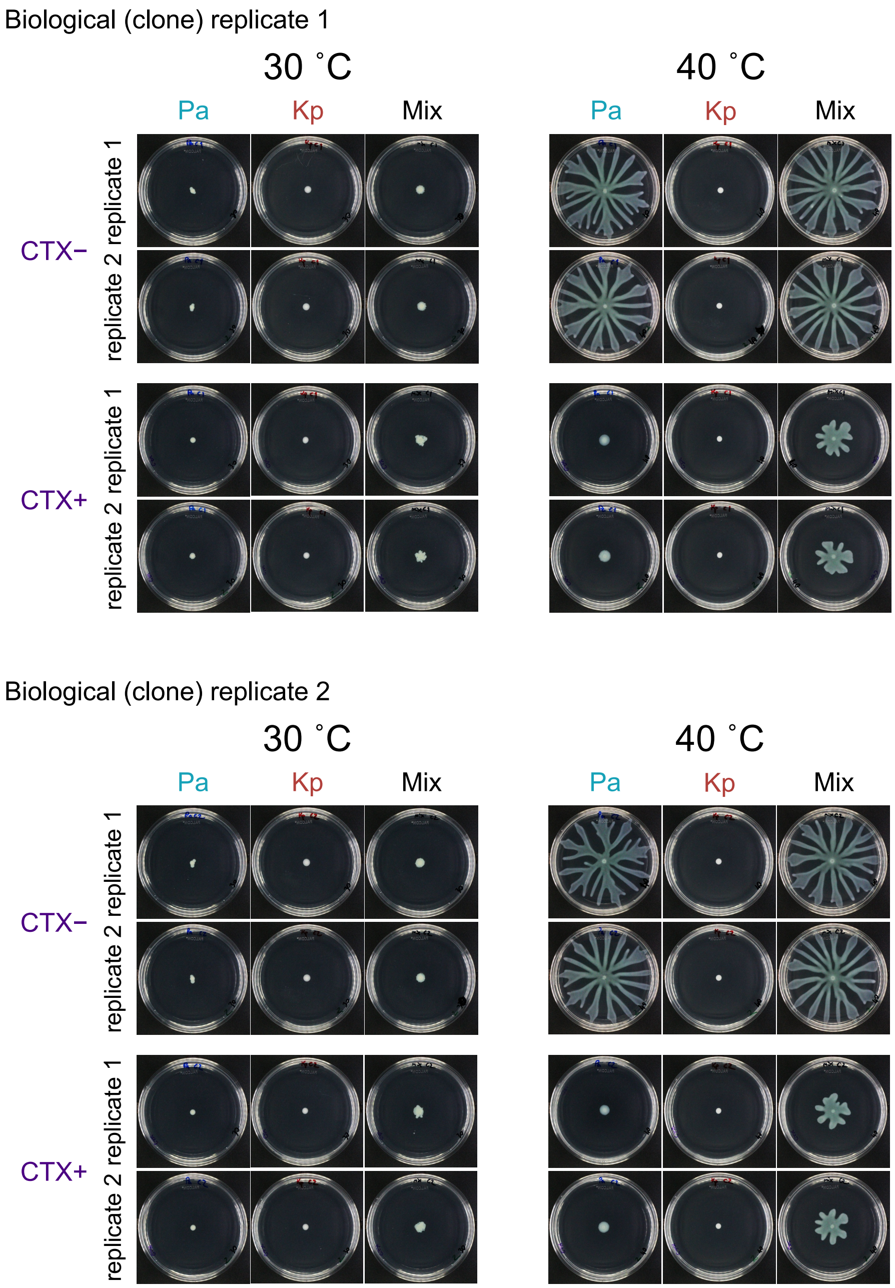


**Supplementary Figure 5: The collective range expansion of *Pseudomonas*-*Klebsiella* under the CTX treatment at different temperature.**

The experiment in Fig. 1c,d was repeated with incubating the agar media at 30 ºC or 40 ºC. For each case, two biological replicates (two different clones) were performed, each with two plate replicates as shown. The images were equally processed for brightness and contrast. CTX (2.5 $\mu$g/ml) was used when applicable.

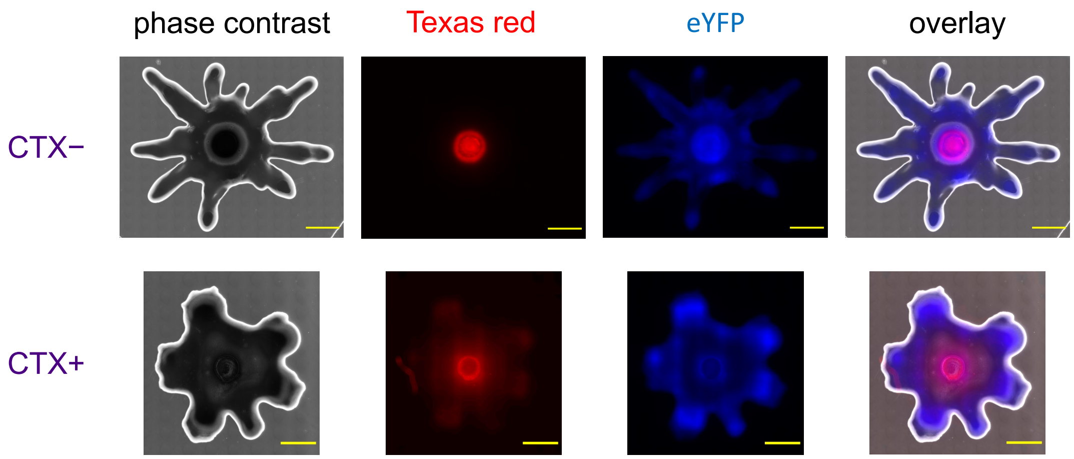


**Supplementary Figure 6: Localizations of *Klebsiella* and *Pseudomonas* in their 'mix' cultures on agar media.**

The cells were prepared as described in Fig. 1. Here, the *Klebsiella* and *Pseudomonas* strains were variants that constitutively expressed mCherry and eYFP, respectively, from plasmids with a gentamycin resistance marker. 20 ml of agar (with casamino acids 10 g/l and agar 0.55 %) media surface was center inoculated with 1 $\mu$l of a 1:1 volume mixture of *Klebsiella* and *Pseudomonas* ($OD_{600}$ = ~ 0.43) without (*top panel*) or with 2.5 $\mu$g/ml of CTX added (*bottom panel*). Gentamycin (10 $\mu$g/ml) and anhydrotetracycline (aTc; 100 ng/ml) were added for plasmid maintenance and mCherry induction, respectively. The images were taken at ~20 h using a three-channel microscopy, phase contrast, Texas red (for *Klebsiella*-mCherry), and eYFP (for *Pseudomonas*-YFP) which were displayed in gray, red, and blue, respectively. A 4× objective was used. The Texas red and eYFP exposure times were 1.2- and 1.5- times greater for the CTX+ case. We ensured no oversaturation of the recorded pixel intensities, allowing us to normalize them during the preparation of the displayed images. The displayed images were downsized 100-times due to large sizes of the original images, which are available upon request. The overlay image was obtained by a manual alignment of the three individual channels. Scale bars: 4 mm.

<u>Image processing protocol:</u>

- Open the stitched images.
- Normalize the pixel intensities for each channel as necessary: For the CTX+ case, divide the pixel intensities by 1.2 and 1.5 for the Texas red and eYFP channels, respectively.

- Scale 0.1 times in both x and y directions.
- Save.
- Crop the images to the rectangle of maximum possible size (1821-by-1473 pixels and 1050-by-1092 pixels for CTX– and CTX+, respectively). Crop the images to ensure the best possible manual alignment of the three channels as the stitched images of the individual channels generally do not come out in equal size.
- Change image types to 16-bit.
- Overlay: Gray for phase contrast, red for Texas red, and blue for eYFP channels.
- Add scale bars to the individual as well as the overlay images (with 15 pixels height).
- Save as png.

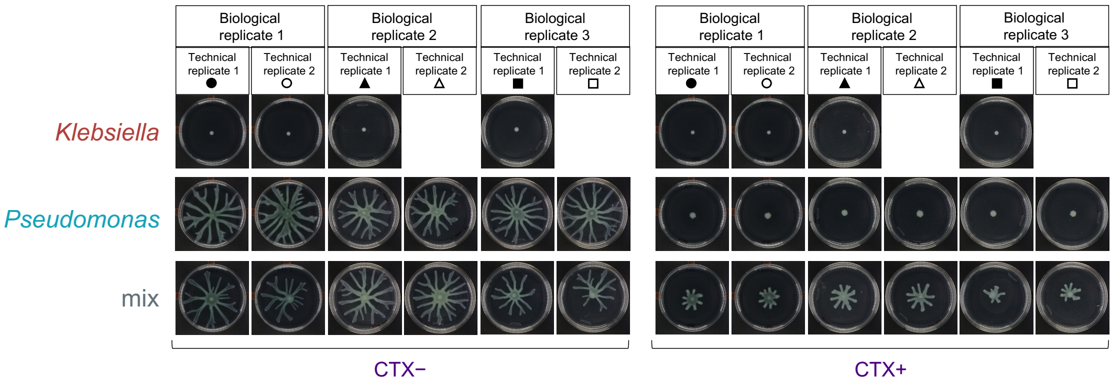


**Supplementary Figure 7: The raw images for Fig. 1c,d.**

The images for Biological replicate 1, Technical replicate 1 were presented in Fig. 1c,d. CTX (2.5 $\mu$g/ml) was used when applicable.

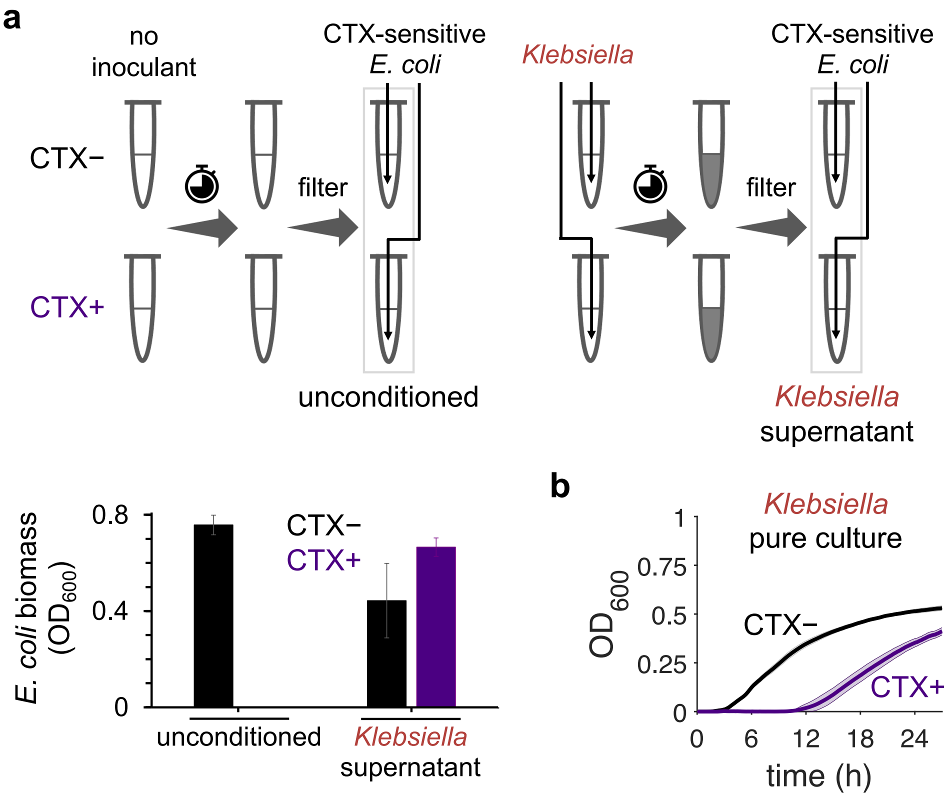


**Supplementary Figure 8: *Klebsiella* degrades CTX in the culture environment.** Cell cultures were prepared as described in Fig. 1.

a. ***Klebsiella* degraded CTX in the culture media.** Culture media were incubated for ~ 6.3 h without or with CTX (2.5 $\mu$g/ml) added and with no inoculant or with *Klebsiella* (initial cell density of $OD_{600}$ = ~ $5\times10^{-4}$). Then, clavulanate (5 $\mu$g/ml) was added to stop Bla activity before supernatants were collected by filtration using 0.22 $\mu$m cellulose acetate filters (VWR). The filtered supernatants were inoculated with CTX-sensitive *E. coli* ($OD_{600}$ = ~ $5\times10^{-4}$). $OD_{600}$ measurements were taken after 21 h of incubation of *E. coli* in the supernatants. All the incubations were performed at 37 °C and 225 r.p.m. Data and error bars represent average and one standard deviation from biological replicates (three and two for the *Klebsiella* supernatant and the unconditioned case, respectively), each with two technical replicates.

b. **Growth curves of *Klebsiella* with the initial cell density used here.** Liquid pure cultures of *Klebsiella* were inoculated, with the initial cell density of $OD_{600}$ = ~ $5\times10^{-4}$, and without (black) or with CTX (2.5 $\mu$g/ml) added (purple). The curves and shades represent the average and the standard deviation, respectively, from two biological replicates each with three technical replicates. See Methods for details.

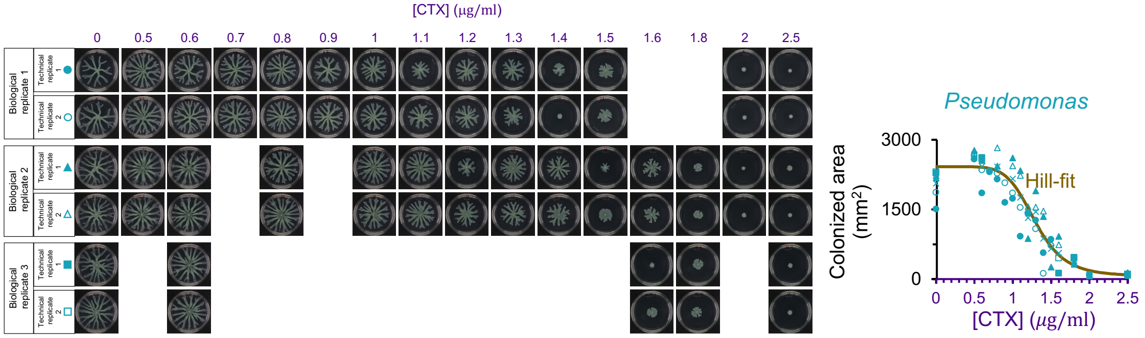


**Supplementary Figure 9: CTX suppresses *Pseudomonas* spatial expansion in a dose-dependent manner.**

*Left panel.* The same spatial expansion experiment in Fig. 1c was repeated with *Pseudomonas* only and varied initial concentration of CTX. The images were taken at 24 h, processed and analyzed for colonized areas using a custom Python script (Methods). Standard 100-mm petri dishes were used.

*Right panel*. The suppression of *Pseudomonas* spatial expansion by CTX fits well to Hill inhibition function (brown, $R^2$= 0.968), given by $2360.3 \times \left(A_C^h/\left(A_C^h + {A_0}^h\right)\right) + 62.3$ mm$^2$, using the built-in lsqcurvefit function of MATLAB 2021b, with $A_C = 1.29\ \mu\mathrm{g/ml}$ and $h = 6.64$ . Circles, triangles, and squares represent biological replicates. Open and solid symbols are two technical replicates. The cross symbols show the average of the replicates.

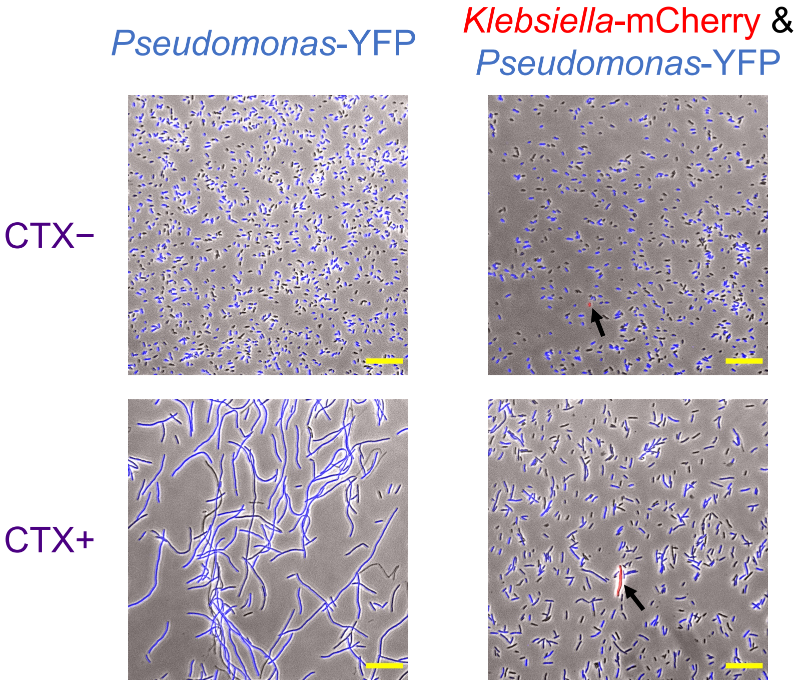


**Supplementary Figure 10: CTX treatment induced *Pseudomonas* cell filamentation, whose extent was reduced in coculture with *Klebsiella*.**

The experiment in Fig. 1c,d was repeated with the strains distinctly labeled by plasmids that constitutively expresses fluorescent proteins, *Pseudomonas*-YFP and *Klebsiella*-mCherry. The cells were harvested at 20 h, suspended in saline, and imaged on coverslips using a three-channel microscopy, eYFP, Texas red (for mCherry), and phase contrast. The images shown are overlays of the three channels after equal adjustment for brightness and contrast, where YFP, mCherry, and phase contrast are displayed in blue, red, and gray, respectively. The coverslips were coated with poly-L-lysine (0.01 %) for enhanced cell immobilization. The black arrows in the right panel point out *Klebsiella*-mCherry cells. CTX (2.5 $\mu$g/ml) was used when applicable. Gentamycin (15 $\mu$g/ml) and aTc (100 ng/ml) were added for plasmid maintenance and mCherry expression, respectively. Scale bars: 20 $\mu$m.

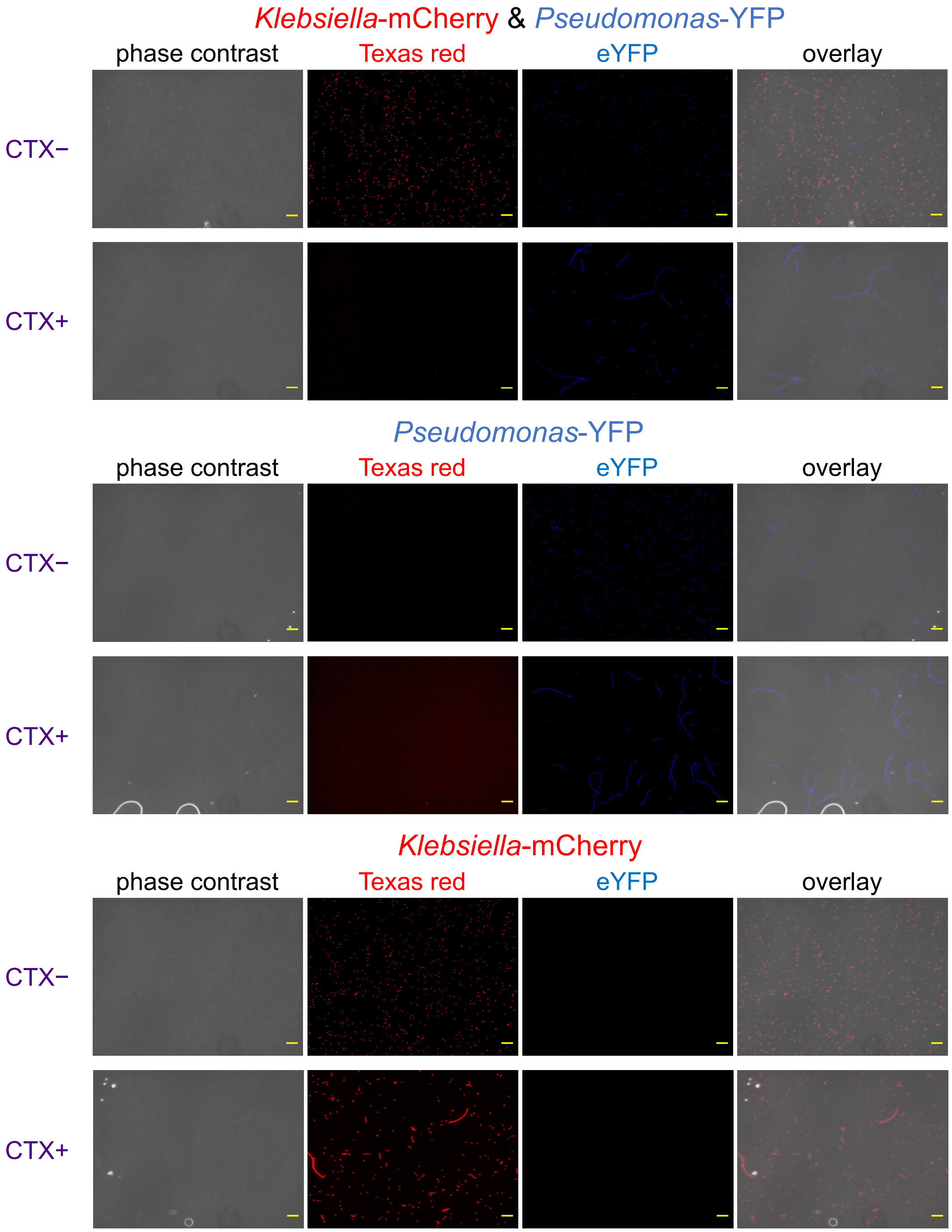


**Supplementary Figure 11: CTX treatment induced *Pseudomonas* cell filamentation in well mixed cultures.**

The experiment in Fig. 1a,b was repeated with the *Pseudomonas*-YFP and *Klebsiella*-mCherry strains. The cells were harvested at 27 h and imaged on coverslips using a three-channel microscopy, eYFP, Texas red (for mCherry), and phase contrast. YFP, mCherry, and phase contrast are displayed in blue, red, and gray, respectively. The resulting images were shown both separately and as an overlay of the three channels after the same adjustments for brightness and contrast. The coverslips were coated with poly-L-lysine (0.01 %) for enhanced cell immobilization. CTX (2.5 $\mu$g/ml) was used when applicable. Gentamycin (15 $\mu$g/ml) and aTc (100 ng/ml) were added for plasmid maintenance and mCherry expression, respectively. Scale bars: 20 $\mu$m.

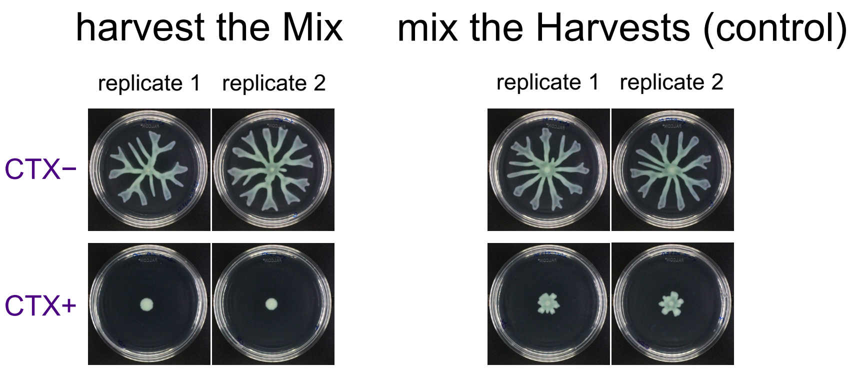


**Supplementary Figure 12: Actively swarming *Pseudomonas* still needed *Klebsiella* for swarming when inoculated on fresh antibiotic media at high cell density.**

As in Fig. 1c,d, we incubated *Pseudomonas* and *Klebsiella* on the swarming agar media as either monocultures without CTX treatment or by first mixing them at 1:1 biomass ratio and with the CTX treatment. During active swarming of the mix culture on the CTX-treated media, we harvested the colony front. In parallel, we also harvested *Pseudomonas* and *Klebsiella* from their respective untreated pure cultures. We adjusted the $OD_{600}$ of these three types of cell harvests to ~2, ~4 times greater than the original inoculum density. Then, we re-inoculated the mix colony harvest ("harvest the Mix" in the figure above) on fresh media with/without the CTX treatment. As a control, we also mixed the *Pseudomonas* and *Klebsiella* pure culture harvests at 1:1 volume ratio ("mix the Harvests" in the figure above) and then inoculated on fresh media with/without the CTX treatment. As a result, the mix culture harvest didn't swarm on the fresh CTX treated media. All other conditions including the control yielded results consistent with our original findings. This suggests that *Klebsiella* was diluted out in the front of the actively swarming mix colony at the harvest time, in agreement with our results in Supplementary Fig. 6. In conclusion, *Klebsiella* is still important for swarming if a swarming activated *Pseudomonas* population is used. The images were taken after 24 h incubation and equally processed for brightness and contrast. CTX (2.5 $\mu$g/ml) was used when applicable.

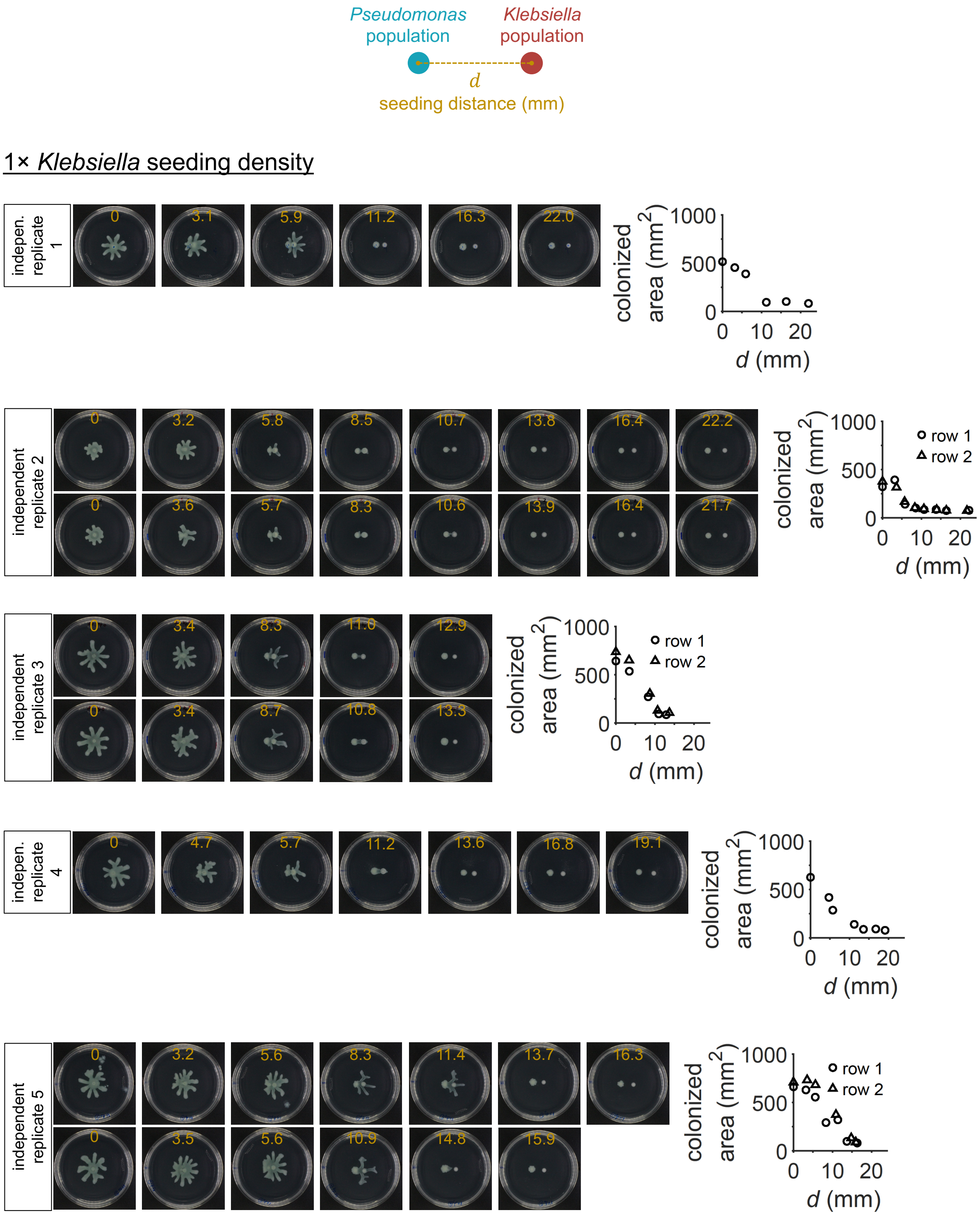

Pseudomonas population
Klebsiella population
d
seeding distance (mm)
1× Klebsiella seeding density
indepen. replicate 1
0
3.1
5.9
11.2
16.3
22.0
colonized area (mm2)
1000
500
0
0 10 20
d (mm)
independent replicate 2
0
3.2
5.8
8.5
10.7
13.8
16.4
22.2
0
3.6
5.7
8.3
10.6
13.9
16.4
21.7
row 1
row 2
independent replicate 3
0
3.4
8.3
11.0
12.9
0
3.4
8.7
10.8
13.3
indepen. replicate 4
0
4.7
5.7
11.2
13.6
16.8
19.1
independent replicate 5
0
3.2
5.6
8.3
11.4
13.7
16.3
0
3.5
5.6
10.9
14.8
15.9

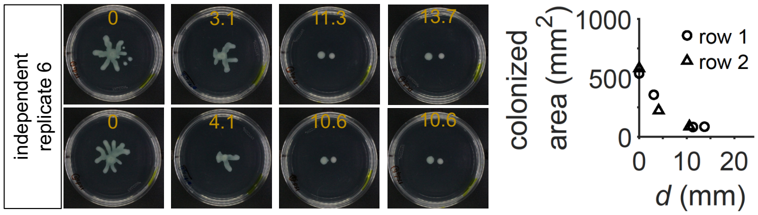


0.1× *Klebsiella* seeding density

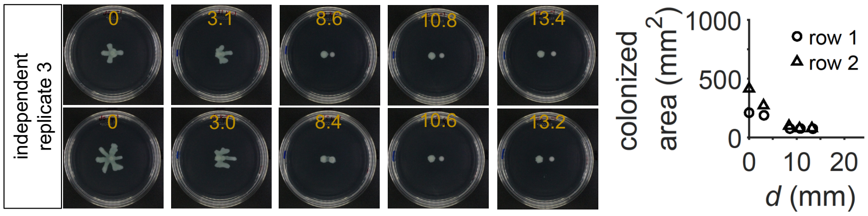


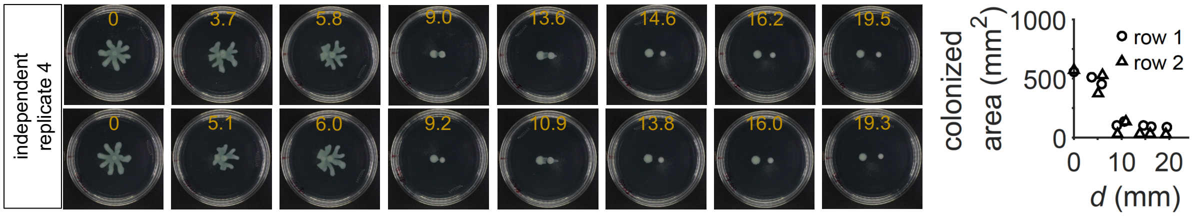


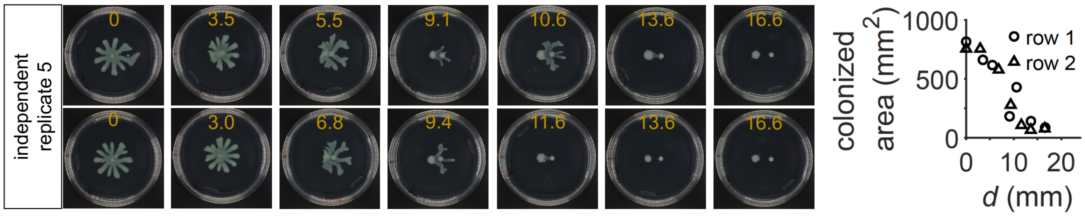


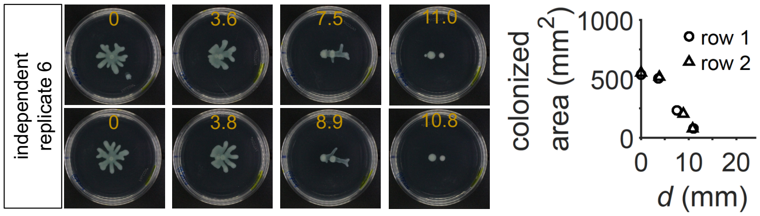


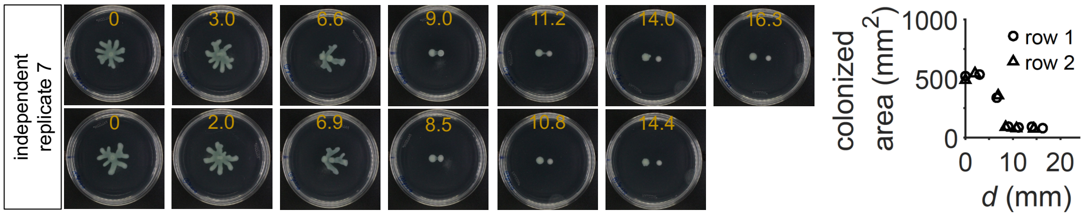

0.5× *Klebsiella* seeding density

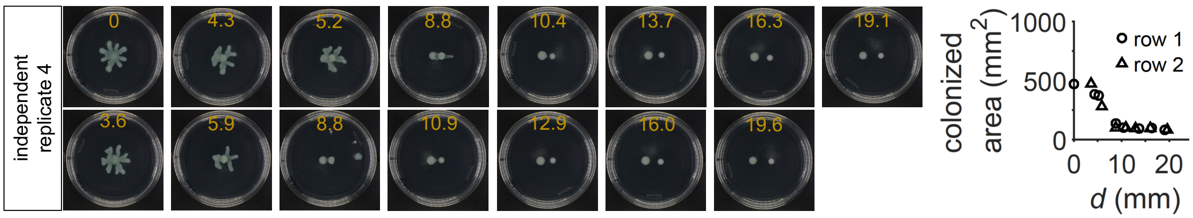


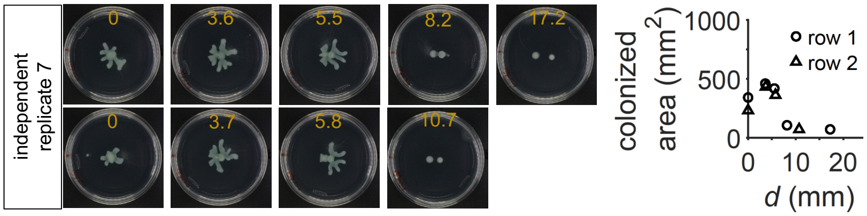


2.5× *Klebsiella* seeding density

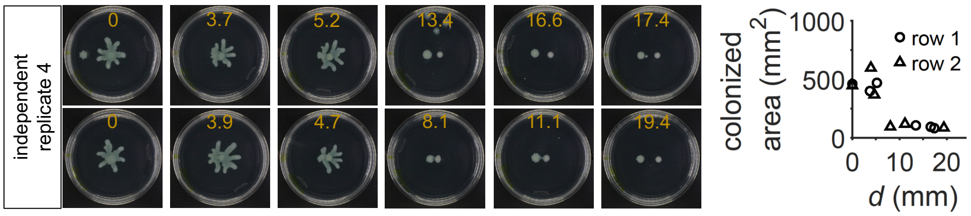


5× *Klebsiella* seeding density

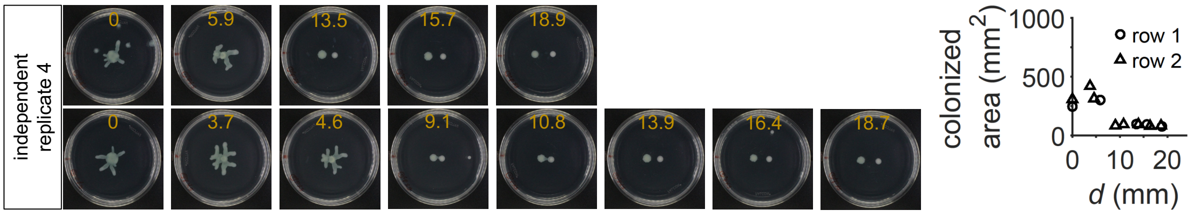


10× *Klebsiella* seeding density

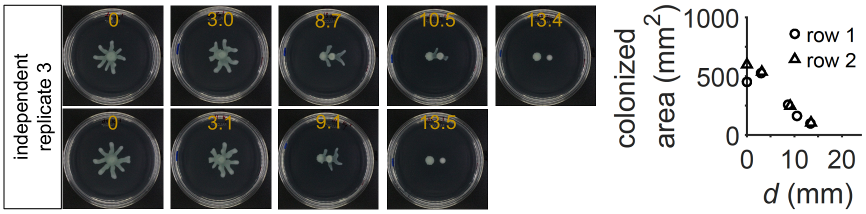

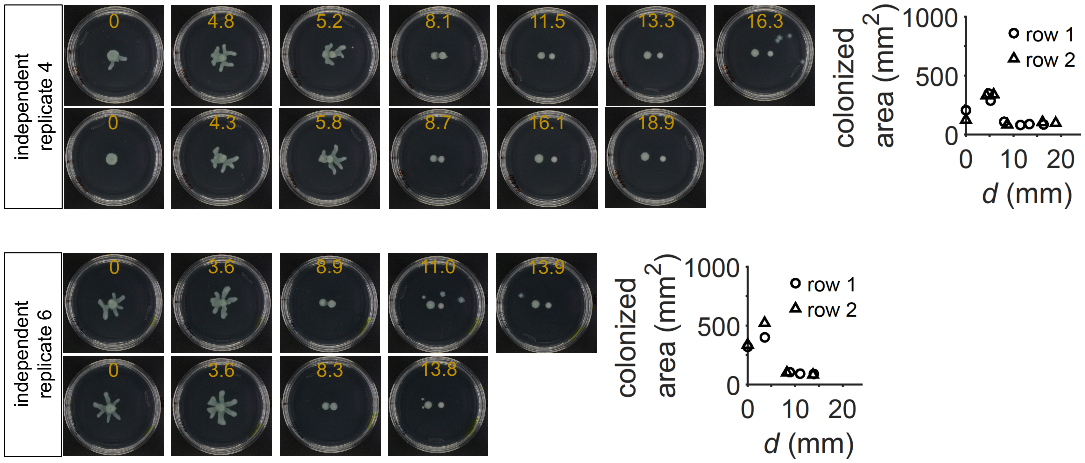


**Supplementary Figure 13: All raw data for Fig. 2b.**

The experiment in Fig. 2b was repeated seven times independently with two technical replica plates when applicable, also in some cases with varied *Klebsiella* seeding cell density. The dishes were imaged at 23 h, and colonized areas were measured. The resultant colonized areas were plotted as a function of the seeding distance (center-to-center) between the *Pseudomonas* and *Klebsiella* populations. The spatial expansion was generally limited to $d$ < 10 mm and similar for different *Klebsiella* seeding cell densities. The data from 1× *Klebsiella* seeding cell density, independent replicate 1 was presented in Fig. 2b.

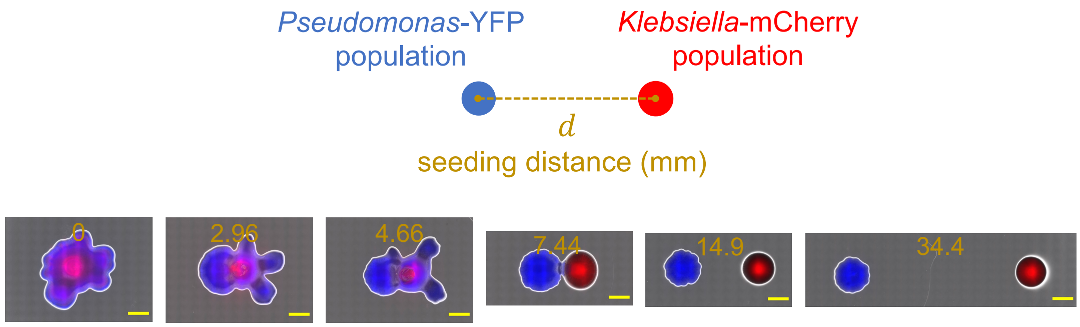


**Supplementary Figure 14: Experiment with the fluorescently labeled *Klebsiella* and *Pseudomonas* strains confirmed the range expansion bias towards *Klebsiella*.**

The experiment in Fig. 2b was repeated using the *Pseudomonas*-YFP and the *Klebsiella*-mCherry variants. The dishes were imaged at 24 h using a Keyence microscope with 4x objective in phase contrast, eYFP (for *Pseudomonas*-YFP), and Texas red (for *Klebsiella*-mCherry) channels. The obtained microscopic images were stitched using a Keyence software. The stitched images were downsized 100-times and overlaid (the original images are available upon request). In the overlay images phase contrast, *Pseudomonas*-YFP, and *Klebsiella*-mCherry are shown in gray, blue, and red, respectively. *Pseudomonas*' spatial expansion is observed for sufficiently small distances (gold in mm) with an evident bias towards the *Klebsiella* population. 0 mm and 2.96 mm Texas ted channel images were obtained using 50- and 7.5-times greater exposure times compared to the rest of the images. Their displayed minimum and maximum intensities were also chosen differently than the others (as 8 and 51, respectively), to remove the background redness. Scale bars: 4 mm.

<u>Image processing protocol:</u>

- Open the stitched images.
- Scale 0.1 times in both x and y directions.
- Save.
- For fluorescence images, change image type → RGB color → 16-bit
- Crop to 1376-by-729 pixels image.
- Overlay
- Crop for display as a 742 by 530 pixels rectangle
- Add 400 $\mu$m (in the downsized image) scale bar with 15 pixels height
- Save as png.

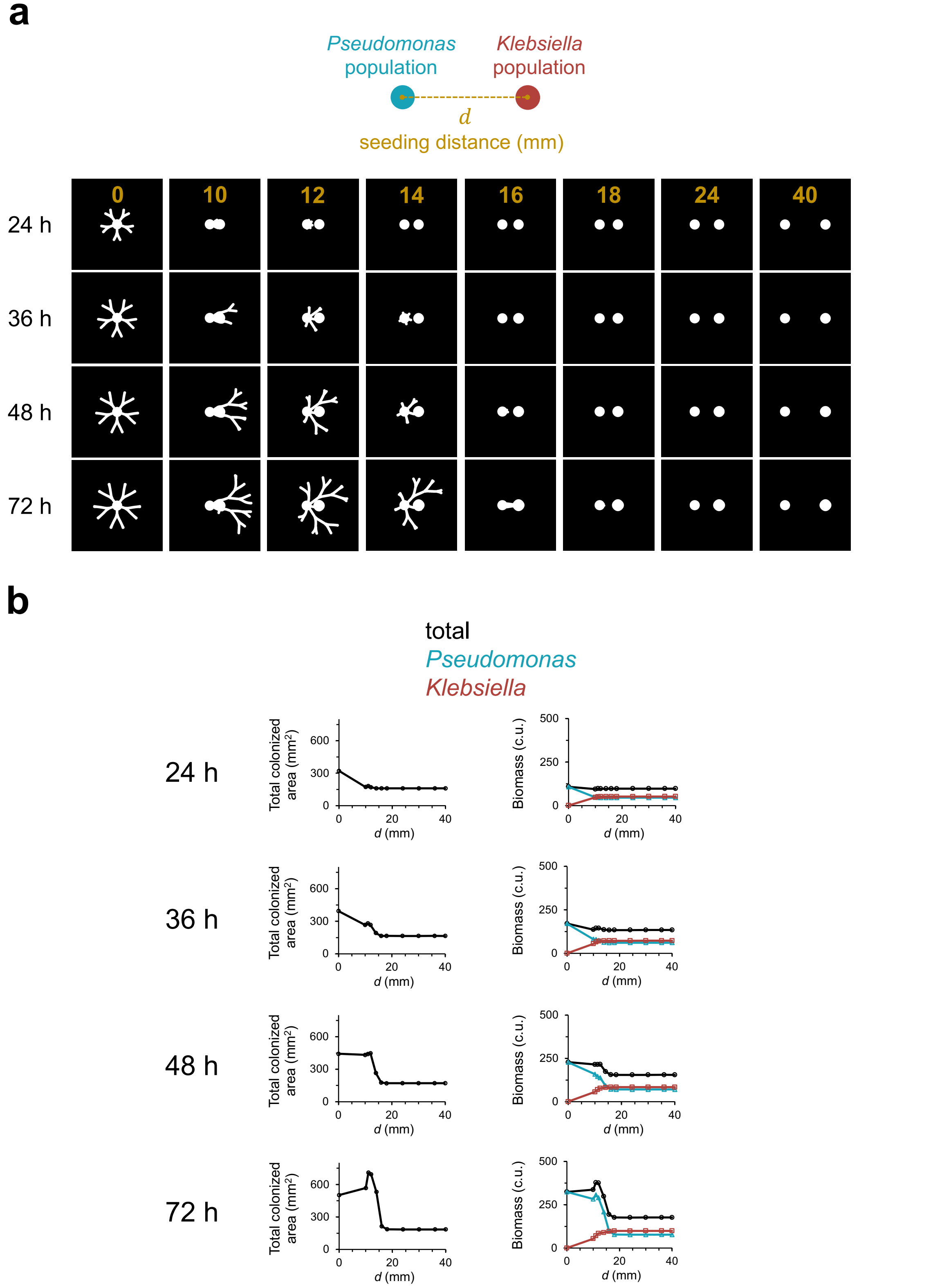


**Supplementary Figure 15: Colonized areas and biomasses at different simulation times.** Data from 48 h were presented in Fig. 2c. Inoculum radius was 5 mm.

**a.** Simulations were run using our mathematical model (Methods). One *Klebsiella* population and one *Pseudomonas* population were inoculated with varied distance $(d)$ between them (gold in mm), as depicted. CTX (2.5 $\mu$g/ml) treatment was mimicked. At each time point shown, matrices representing spatial distributions of *Pseudomonas* and *Klebsiella* cell densities were overlaid. The overlay image was converted to black-white image using imbinarize function of MATLAB 2021b with thresholding for the initial local cell density value.

**b.** For each case in (a), the total number of occupied pixels were calculated and multiplied by 0.0081 $mm^2$, the pixel to physical area conversion factor for our simulation domains. The resulting colonized areas are shown alongside the total, *Pseudomonas*, or *Klebsiella* biomasses.

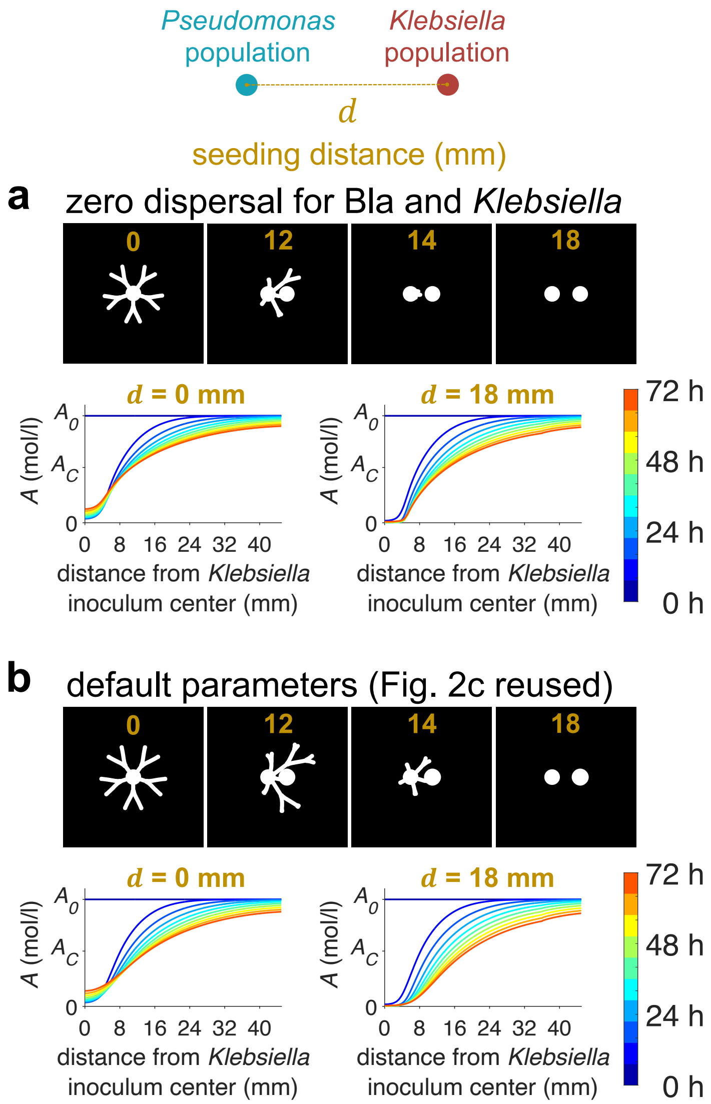


**Supplementary Figure 16: Simulations mimicking an absolutely local, intracellular-only degradation of antibiotics.**

**a.** *Top panel*. Coarse-grained kinetic model used in Fig. 2c was simulated using zero dispersal for Bla and *Klebsiella* that produces it ($D_B = 0$ and $D_E = 0$). This effectively represents a system in which antibiotic degradation is by an immotile species and intracellular-only. Results shown are for *t* = 48 h. *Bottom panel*. For $d = 0$ mm and $d = 18$ mm, as representatives of the maximum and zero spatial expansion abilities respectively, azimuthally averaged profile of the antibiotic concentration (*A*) with respect to the *Klebsiella* inoculum center was computed for at every eight hour of a 72-h simulation.

**b.** The data in Fig. 2c was reused (*top*) and the same azimuthally averaged *A* computations were performed (*bottom*), for comparison.

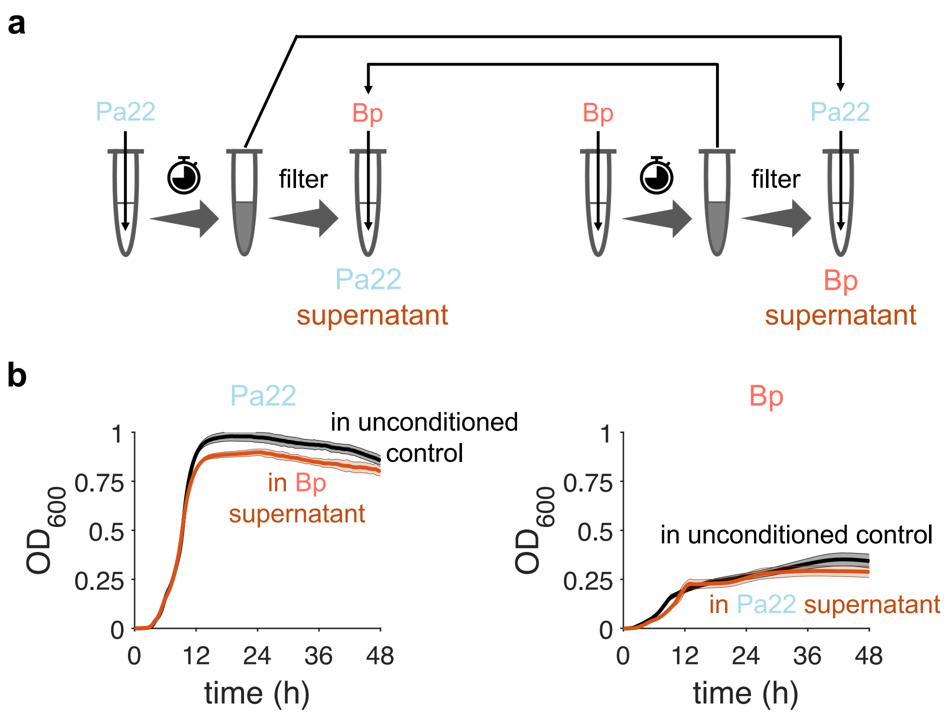


**Supplementary Figure 17: Spent media experiments demonstrate that Pa22 and Bp primarily compete.**

**a.** Culture media were incubated for ~ 6.3 h at 37 °C and 225 r.p.m. with no inoculant (for an unconditioned control media) or with Bp or Pa22 (initial cell density of $OD_{600}$ = ~ $5\times10^{-4}$). Then, supernatants were collected by filtration using 0.22 $\mu$m cellulose acetate filters (VWR). The filtered supernatants of each species were cross-inoculated with the other species ($OD_{600}$ = ~ $5\times10^{-4}$).

**b.** $OD_{600}$ measurements were taken in a plate reader at 37 °C. For each case, six replicates were performed—two biological replicates (two different clones) each with three technical replicates (wells) for the growth in the supernatants. The curves and shades represent the average and the standard deviation, respectively, from all replicates.

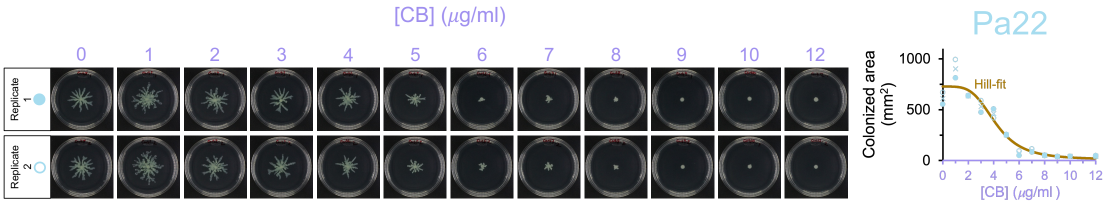


**Supplementary Figure 18: CB inhibits Pa22's spatial expansion in a dose-dependent manner.**

*Left panel.* The same experiment in Fig. 3c was repeated with Pa22 only and varied initial concentrations of CB. The images were taken at 24 h, processed and analyzed for colonized areas using a custom Python script (Methods). Standard 100-mm petri dishes were used.

*Right panel*. The suppression of Pa22's spatial expansion by CB fits well to Hill inhibition function (brown, $R^2$= 0.940), given by $717.4 \times \left(A_C^h/\left(A_C^h + {A_0}^h\right)\right) + 10.2$ mm$^2$, using the built-in lsqcurvefit function of MATLAB 2021b. $A_0$ is the initial concentration of CB, $A_C = 4.145\ \mu\text{g/ml}$ is the half-maximum suppressing value of $A_0$, and $h = 4.02$ is the Hill coefficient ($R^2$= 0.9401). Open and solid symbols are two technical replicates. The cross symbols show the average of the replicates.

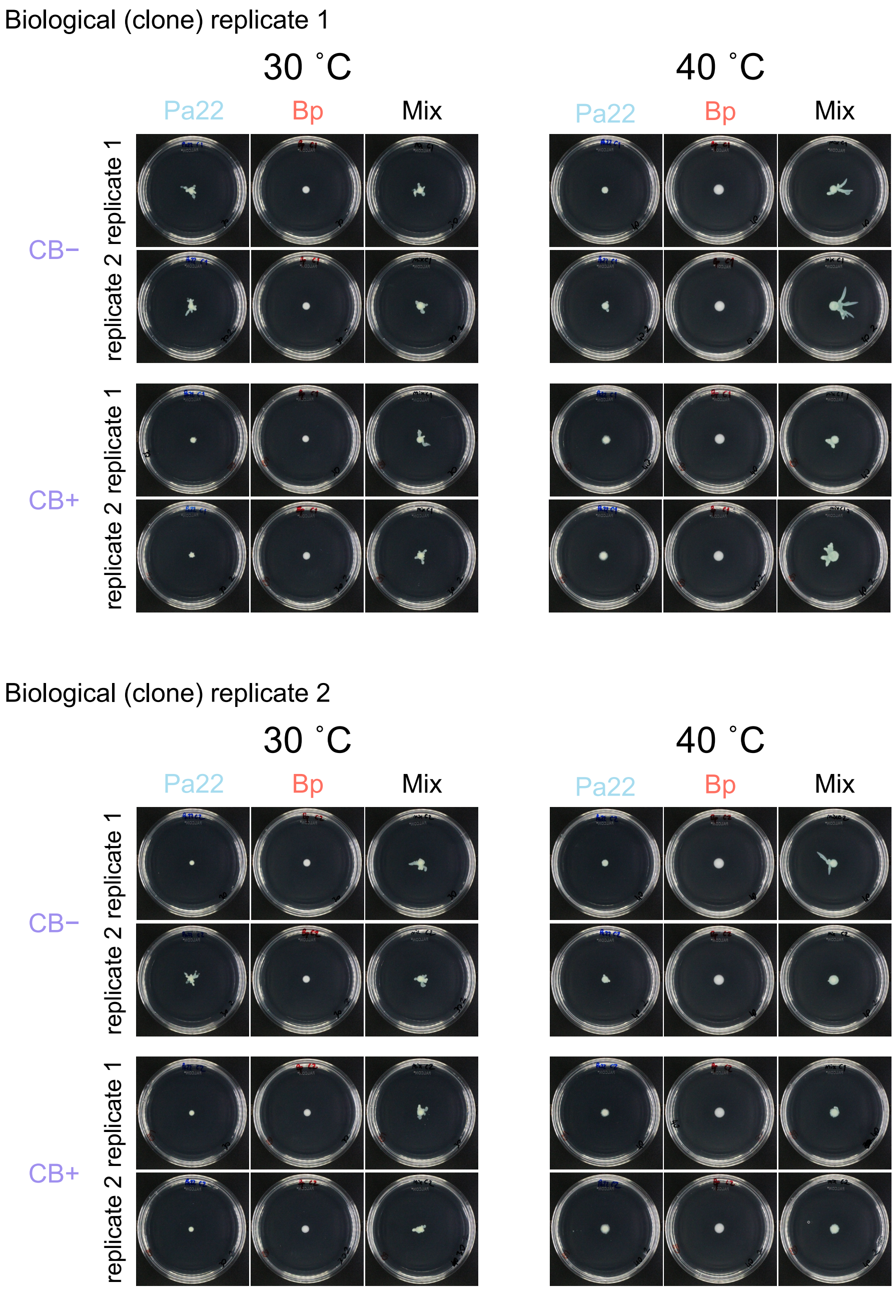


**Supplementary Figure 19: The collective range expansion of Pa22 and Bp under the CB treatment at different temperatures.**

The experiment in Fig. 3c,d was repeated with incubating the agar media at 30 ºC or 40 ºC. For each case, two biological replicates (two different clones) were performed, each with two plate replicates as shown. The images were equally processed for brightness and contrast. CB (10 $\mu$g/ml) was used when applicable.

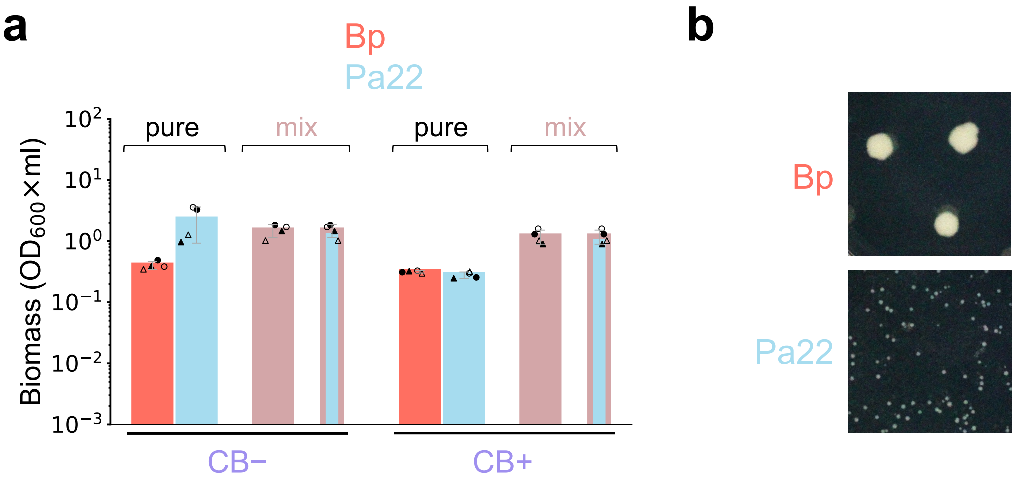


**Supplementary Figure 20: Bp and Pa22 biomass production in pure- and mix-cultures on agar, in which the latter was distributed based on the distinct colony forming unit (CFU) morphologies of the two species.**

**a.** The total biomass of the cell harvests in Fig. 3c,d was calculated by measuring the absorbance ($OD_{600}$) of the harvest suspension and multiplying it by the harvest's volume. Relative contributions of Bp (red) and Pa22 (light blue) to the total biomass values for "mix" (the colored bars with a pink frame) were estimated by multiplying the total "mix" biomass with the relative abundances of Bp and Pa22 (Fig. 3d), determined by plating based on distinct CFU morphologies of the two species. Circles (inoculum density of $OD_{600}$ = ~ 0.43) and triangles (inoculum density of $OD_{600}$ = ~ 0.34) biological replicates whose technical replicates were shown by solid or open markers. The data and error bars represent the mean and standard deviation, respectively, from the biological replicates after their technical replicates were averaged.

**b.** **Distinct CFU morphologies of Bp and Pa22.** We distinguished between Bp and Pa22 by plating them on LB agar (1.5 %) plates and categorizing them the following day based on their distinct CFU appearances. Here we show example images for Bp (top) and *Pseudomonas* (bottom) colonies in the same scale.

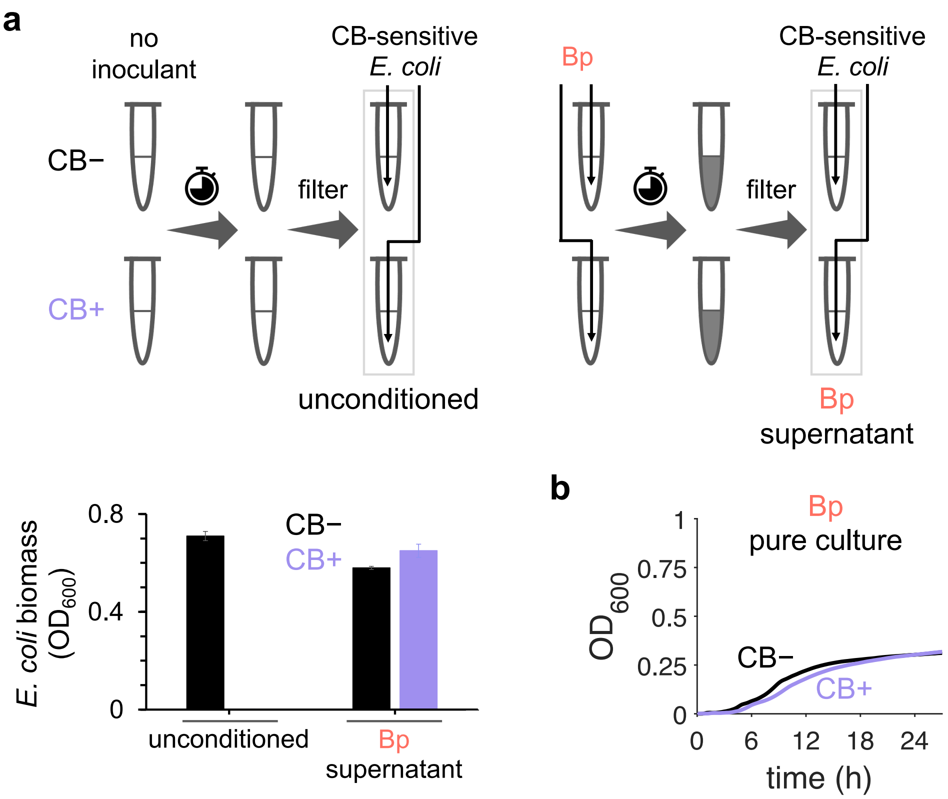


**Supplementary Figure 21: Bp degrades CB in the culture environment.** Cell cultures were prepared as described in Fig. 3.

a. **Bp degraded CB in the culture media.** Culture media were incubated without or with CB (10 $\mu$g/ml) added, and with no inoculant or with Bp (initial cell densitiy of $OD_{600}$ = ~ $5\times10^{-4}$) for ~ 6.3 h. Then, clavulanate (5 $\mu$g/ml) was added to stop any potential Bla activity before supernatants were collected by filtration using 0.22 $\mu$m cellulose acetate filters (VWR). The filtered supernatants were inoculated with CB-sensitive *E. coli* ($OD_{600}$ = ~ $5\times10^{-4}$). $OD_{600}$ measurements were taken after 21 h of incubation of *E. coli* in the supernatants. All the incubations were performed at 37 °C and 225 r.p.m. Data and error bars represent average and one standard deviation from three biological replicates, each with two technical replicates.

b. **Growth curves of Bp with the initial cell density used here.** Liquid pure cultures of Bp were inoculated, with the initial cell density of $OD_{600}$ = ~ $5\times10^{-4}$, and without (black) or with CB (10 $\mu$g/ml) added (violet). The curves and shades represent the average and the standard deviation, respectively, from two biological replicates each with three technical replicates. See Methods for details.

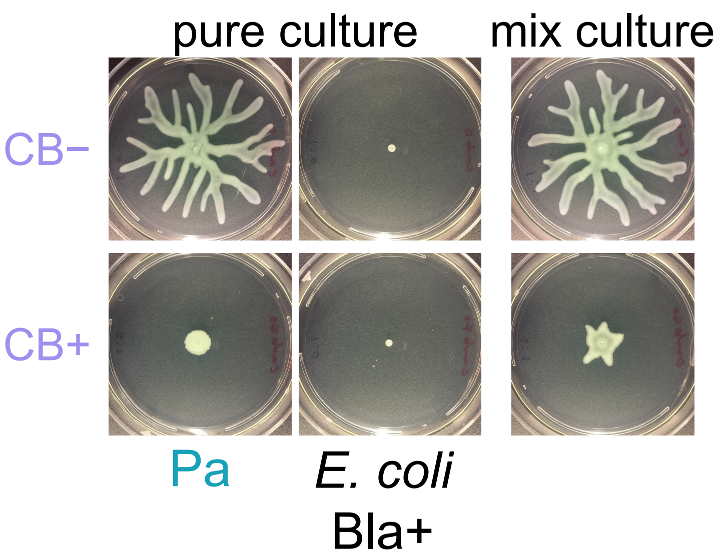


**Supplementary Figure 22: The collective range expansion is generalizable.**

We used an *E. coli* strain expressing Bla (*E. coli* Bla+) from a plasmid conferring carbenicillin resistance. Cultures of single clones of this *E. coli* Bla+ and *Pseudomonas* (Pa) were separately grown overnight in LB. 20 ml of agar media (phosphate-buffered saline supplemented with 8 g/l of casamino acids and agar 0.5%) without or with CB (40 $\mu$g/ml) was center inoculated with 1 $\mu$l of *E. coli* Bla+ and Pa as pure cultures or a 1:1 volume mixture (mix culture). The images of the colonies were captured at 20 h. Standard 100-mm petri dishes were used. Isopropyl β-D-1-thiogalactopyranoside (IPTG) was added at 1 mM final concentration for inducing the Bla expression.

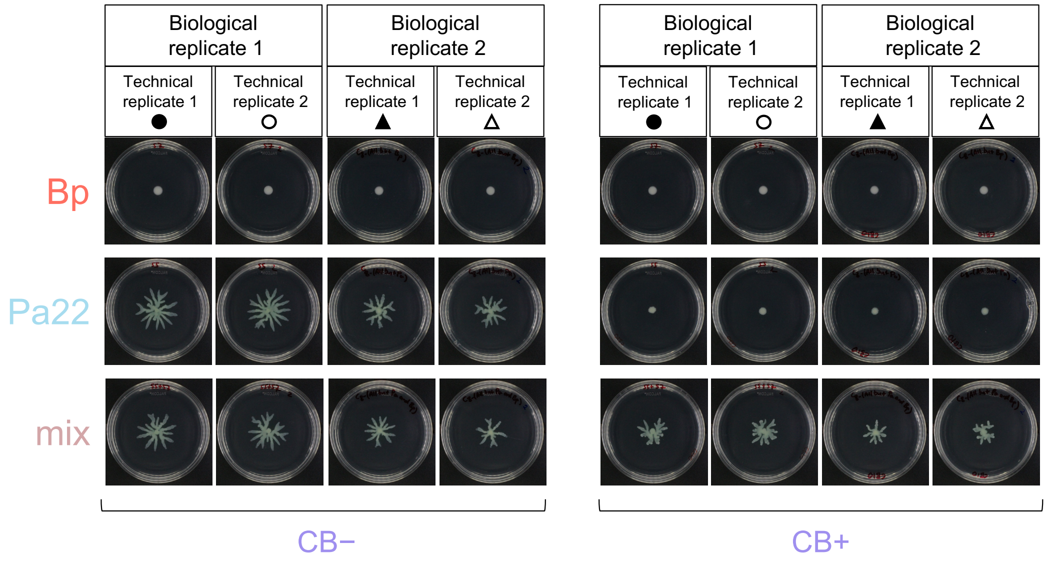


**Supplementary Figure 23: The raw images for Fig. 3c,d.**

The images for Biological replicate 1, Technical replicate 2 were presented in Fig. 3c,d. CB (10 $\mu$g/ml) was added when applicable. Biological replicates 1 and 2 had inoculum cell densities of $OD_{600}$ = ~ 0.43 and $OD_{600}$ = ~ 0.34, respectively.

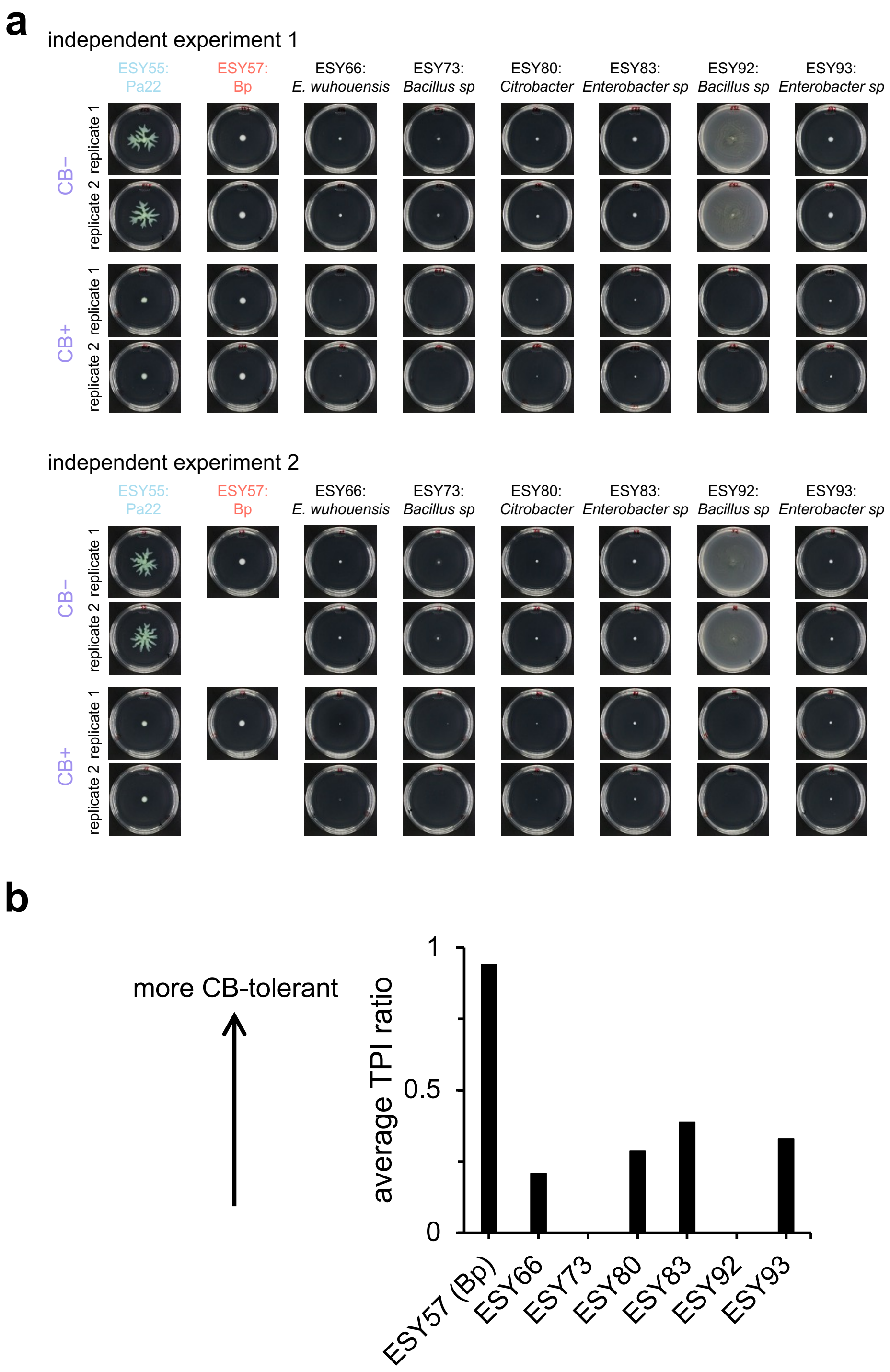


TPI: total pixel intensity

**Supplementary Figure 24: Spatial expansion abilities and CB tolerances of the SynkC members.**

In Fig. 4, an 8-member synthetic community of sink isolates, which we coined 'SynkC', was used. SynkC was assembled by mixing six additional co-isolates with Bp and Pa22. 16S rDNA sequencing categorized the six isolates into four additional, distinct taxonomic groups.

- **a.** The experiment in Fig3c was repeated for each SynkC member. The images were equally processed for their brightness and contrast for presentation. CB (10 $\mu$g/ml) was used when applicable.
- **b.** We analyzed the images in (a) and obtained a quantitative measurement of the CB tolerances of the strains. Using ImageJ, we first subtracted the background intensities from the images. Then, we calculated the total pixel intensity (TPI) for each strain with or without the CB treatment (CB+ and CB–, respectively) and estimated a TPI ratio by dividing the value for CB+ by the value for CB–. The data bars represent the averages from all replicate images shown in (a). Pa22 and ESY92 were excluded from this analysis because their spatial expansions could confound the interpretation of the results here.

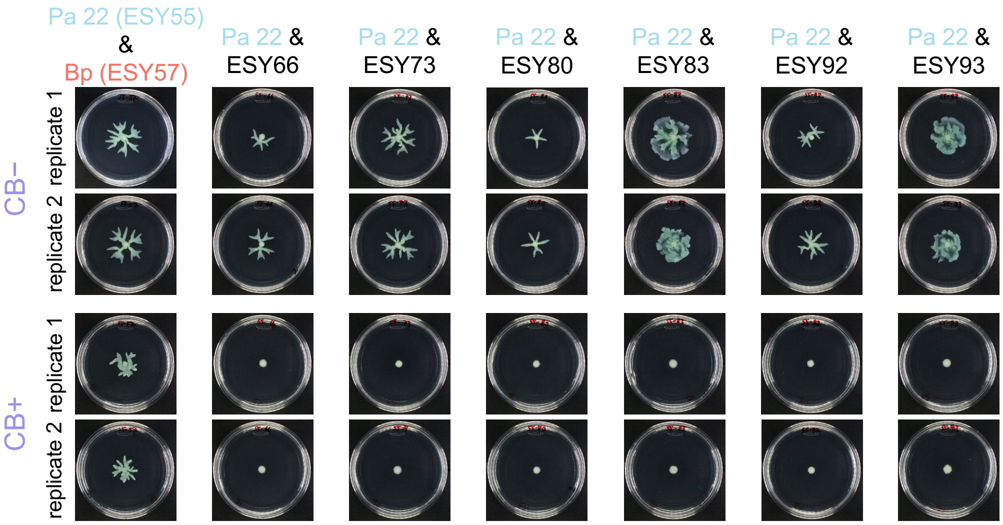


**Supplementary Figure 25: Bp is the only SynkC member enabling collective range expansion of Pa22 under the CB treatment in pairwise cultures.** The experiment was set up as described for Fig. 3d and, in this case, each of the seven pairwise cultures with Pa22 was tested. The images were taken after 24 h and equally processed for their brightness and contrast for presentation.

*Independent replicate 1, Technical replicate 1*

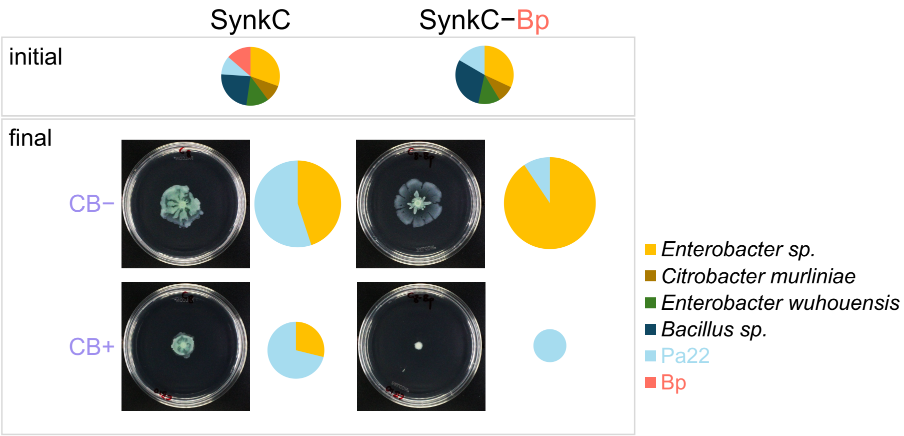


*Independent replicate 1, Technical replicate 2*

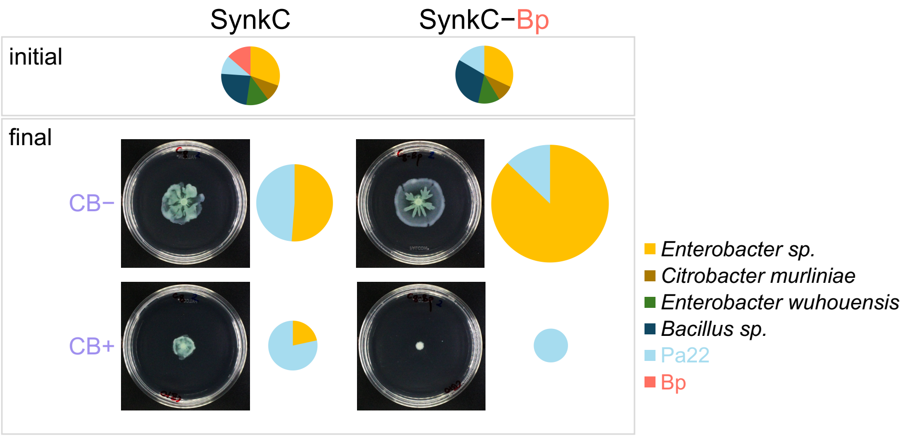


*Independent replicate 2, Technical replicate 1*

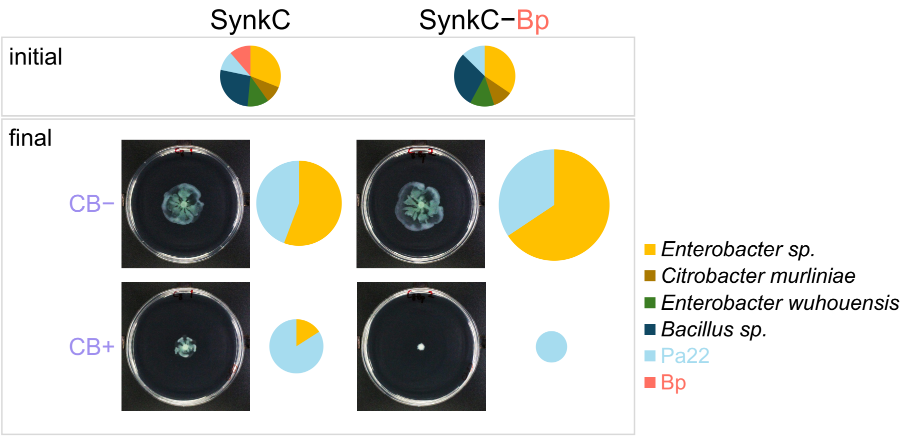

*Independent replicate 2, Technical replicate 2*

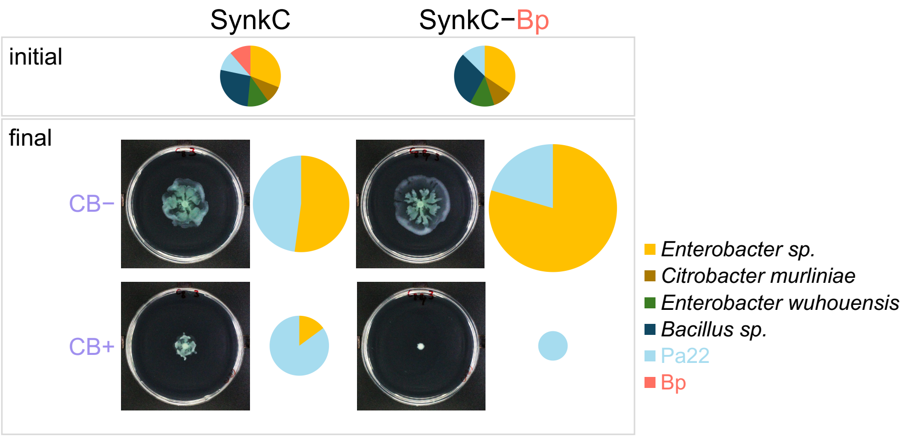


**Supplementary Fig. 26: All the data for Fig. 4.**

The raw images and the relative abundance datasets for all replicates supporting Fig. 4 are presented above. Each independent replicate had two technical replicates, which were inoculated using the same set of initial cell suspensions and, thus, displayed using a single set of pie charts for their initial state above. As in Fig. 4, the community compositions were determined by 16S microbiome sequencing and presented by applying a 0.5% cut-off value. For each independent experiment, the images were processed equally for their brightness and contrast for presentation. Each of the pie charts above shows six groups, with the *Enterobacter sp*. and *Bacillus sp*. groups representing the combined abundance of two isolates each, while all other groups represent a single isolate. For the final time point, the relative areas represent the distribution of the respective biomasses measured, across all pie charts. The ‘Independent replicate 1, Technical replicate 2’ dataset was presented in Fig. 4. CB (10 $\mu$g/ml) was used when applicable.

**Supplementary Table 1: Bla genes found in the strains used in this study.**

| Strain ID | Strain name | Bla gene | Protein product | Bakta annotated locus tag |
|---|---|---|---|---|
| ESY22 | *Pseudomonas* | *ampC* | Beta-lactamase class C family | OFHKIH_07310 |
| | | *ampC* | Beta-lactamase class C family | OFHKIH_09200 |
| | | *blaOXA-488* | OXA-50 family oxacillin-hydrolyzing class D beta-lactamase OXA-488 | OFHKIH_09340 |
| | | *blaPDC-34* | Class C beta-lactamase PDC-34 | OFHKIH_10585 |
| | | *ampC* | Beta-lactamase class C family | OFHKIH_11185 |
| | | *ampC* | Beta-lactamase class C family | OFHKIH_16520 |
| | | *ampC* | Beta-lactamase class C family | OFHKIH_28845 |
| ESY23 | *Klebsiella* | *ampC* | Beta-lactamase class C family | AILCJE_05920 |
| | | *blaLEN* | LEN family class A beta-lactamase | AILCJE_12825 |
| | | *blaCTX-M-9* | Extended-spectrum class A beta-lactamase CTX-M-9 | AILCJE_28575 |
| ESY55 | Pa22 | *ampC* | Beta-lactamase class C family | PINPMI_00865 |
| | | *ampC* | Beta-lactamase class C family | PINPMI_04005 |
| | | *ampC* | Beta-lactamase class C family | PINPMI_04495 |
| | | *blaOXA-50* | Oxacillin-hydrolyzing class D beta-lactamase OXA-50 | PINPMI_04635 |
| | | *ampC* | Beta-lactamase class C family | PINPMI_10820 |
| | | *blaPDC-8* | Class C beta-lactamase PDC-8 | PINPMI_27025 |
| | | *ampC* | Beta-lactamase class C family | PINPMI_28915 |
| ESY57 | Bp | *ampC* | Beta-lactamase class C family | HHBGHA_03180 |
| | | *penP* | Beta-lactamase class A | HHBGHA_05940 |
| | | *bla* | Class A beta-lactamase Bla1 | HHBGHA_06980 |
| | | *ampC* | Beta-lactamase class C family | HHBGHA_07930 |
| | | *ampC* | Beta-lactamase class C family | HHBGHA_08065 |

| | | | | |
|---|---|---|---|---|
| | | *ampC* | Beta-lactamase class C family | HHBGHA_08415 |
| | | *ampC* | Beta-lactamase class C family | HHBGHA_09680 |
| | | *ampC* | Beta-lactamase class C family | HHBGHA_10140 |
| | | *ampC* | Beta-lactamase class C family | HHBGHA_10420 |
| | | *ampC* | Beta-lactamase class C family | HHBGHA_10425 |
| | | *bla* | Class A beta-lactamase | HHBGHA_10510 |
| | | *ampC* | Beta-lactamase class C family | HHBGHA_10760 |
| | | *bla2* | BcII family subclass B1 metallo-beta-lactamase | HHBGHA_11440 |
| | | *blaIII* | Class A beta-lactamase BlaIII | HHBGHA_15295 |
| | | *ampC* | Beta-lactamase class C family | HHBGHA_18510 |
| | | *ampC* | Beta-lactamase class C family | HHBGHA_23245 |
| | | *ampC* | Beta-lactamase class C family | HHBGHA_27625 |
| ESY66 | *Enterobacter wuhouensis* | *blaACT-64* | Cephalosporin-hydrolyzing class C beta-lactamase ACT-64 | IJHIMG_09800 |
| ESY73 | *Bacillus sp.* | *blaBPU* | BPU family class D beta-lactamase | DIBKGJ_12165 |
| ESY80 | *Citrobacter murliniae* | *blaCMY* | CMY-2 family class C beta-lactamase | LOHOIH_04600 |
| ESY83 | *Enterobacter sp.* | *blaACT-64* | Cephalosporin-hydrolyzing class C beta-lactamase ACT-64 | CMIIBO_15550 |
| ESY92 | *Bacillus sp.* | *blaBPU* | BPU family class D beta-lactamase | LPIEEI_12160 |
| ESY93 | *Enterobacter sp.* | *blaACT-64* | ACT family cephalosporin-hydrolyzing class C beta-lactamase | DJFABJ_14990 |

Notes:

1. The table excludes genes for proteins with structural similarities with Bla enzymes but with no known direct evidence in antibiotic resistance (e.g., *gloB*, *ulaG*, *hcpC*, and *romA* as well as those identified as “n/a” and annotated as “beta-lactamase”).

2. All genes annotated as ‘*ampC*’ are reported, regardless of the strength of evidence for their Bla activity.

**Supplementary Table 2: Coarse-grained kinetic model parameters and initial conditions.**

| Parameter | Definition | Value | Units | Reference |
|---|---|---|---|---|
| $D_N$ | Diffusivity of casamino acids | 5.7486 | $mm^2/h$ | 1 |
| $\alpha_P$ | Growth rate constant of *Pseudomonas* | 1.105 | 1/h | 1 |
| $K_N$ | Casamino acids concentration for half-maximum growth rate of *Pseudomonas* | 0.6635 | g/l | 1 |
| $C_m$ | Half-maximum growth rate parameter | 0.0789 | cell density unit, cu. ($OD_{600}$) | 1 |
| $K_C$ | Local carrying capacity | 1 | cell density unit, cu. ($OD_{600}$) | |
| $\beta_N$ | Nutrient consumption rate constant by *Pseudomonas* cell growth | 195.4832 | g/(l.h.cu.) | 1 |
| $\gamma$ | Pseudomonas spatial expansion efficiency | 5 | mm/(h.cu.) | 1 |
| $W$ | Branch width | 2.625 | mm | 1 |
| $D$ | Branching density | 0.1513 | 1/mm | 1 |
| $\kappa_B$ | Rate constant for cefotaxime degradation by Bla | $10^{10}$ | l/(mol. h) | 2 |
| $d_B$ | Spontaneous degradation rate constant for Bla | 0.02 | 1/h | 2 |
| $\phi_E$ | Proportionality between growth and lysis rates of *Klebsiella* | 4 | 1 | 2 |
| $A_E$ | Cefotaxime concentration for half-maximum *Klebsiella* lysis rate | $2\times10^{-6}$ | mol/l | 2 |
| $B_{in}$ | Intracellular concentration of Bla (for *Klebsiella* only) | $3\times10^{-19}$ | mol/cell | 2 |
| $h_2$ | Steepness of the onset of lysis mediated by cefotaxime | 3 | 1 | 2 |
| $c_F$ | $OD_{600}$ to cell/l conversion factor | $1\times10^{12}$ | cell/l | |
| $\beta_{EN}$ | Nutrient consumption rate constant by *Klebsiella* cell growth | 9.77416 | g/(l.h.cu.) | |
| $\mu_E$ | Growth rate proportionality of *Klebsiella* relative to *Pseudomona*s | 1.2 | 1 | |
| $A_C$ | Cefotaxime concentration for half-maximum suppression of *Pseudomonas'* effective spatial expansion ability | $2.7080\times10^{-6}$ | mol/l | |
| $h$ | Steepness of the suppression of *Pseudomonas*' effective spatial expansion ability by cefotaxime | 6.642 | 1 | |
| $D_A$ | Diffusivity of cefotaxime | 6.5 | $mm^2/h$ | |
| $D_B$ | Diffusivity of Bla | 0.114972 | $mm^2/h$ | |
| $D_E$ | Diffusive dispersal rate of *Klebsiella* | $1\times10^{-6}$ | $mm^2/h$ | |

| Initial conditions | | | | |
|---|---|---|---|---|
| $P(t=0)$ | *Pseudomonas* seeding cell density | 0.05 | cell density unit, cu. ($OD_{600}$) | |
| $E(t=0)$ | *Klebsiella* seeding cell density | 0.05 | cell density unit, cu. ($OD_{600}$) | |
| $N(t=0)$ | Initial casamino acids concentration | 14.5 | g/l | 1 |
| $A(t=0)$ | Initial cefotaxime concentration | $5.24\times10^{-6}$ | mol/l | |
| $B(t=0)$ | Extracellular Bla concentration | 0 | | 2 |

[1] Luo, N., Wang, S., Lu, J., Ouyang, X. & You, L. Collective colony growth is optimized by branching pattern formation in *Pseudomonas aeruginosa*. *Mol Syst Biol* **17**, e10089, doi:https://doi.org/10.15252/msb.202010089 (2021).

[2] Meredith, H. R. et al. Applying ecological resistance and resilience to dissect bacterial antibiotic responses. *Science Advances* 4, eaau1873, doi:doi:10.1126/sciadv.aau1873 (2018).

**Supplementary Table 3: Primers used in this study.**

| **Amplified region** | **Direction** | **Sequence (5'→ 3')** |
|---|---|---|
| ptetmCherry backbone | Forward | TTTAGCTTCCTTAGCTCC |
| ptetmCherry backbone | Reverse | TCGACCCAAGTACCGCCACCTAATTTGATATCGAGCTCGC |
| Gentamicin resistance gene | Forward | CAGGAGCTAAGGAAGCTAAAATGTTACGCAGCAGCAAC |
| Gentamicin resistance gene | Reverse | GGTGGCGGTACTTGG |
| Transcriptional terminator | Forward | CGCTTAATTAATTAATCTAGTTCAGCCAAAAAACTTAAGACCG |
| Transcriptional terminator | Reverse | GGACCAAAACGAAAAAAGGC |

### Supplementary Information

Here, we derive analytical solutions to develop intuition on a few critical quantities:

1. **Bla diffusion length scale (*L*)**

   a. <u>Negligible Bla degradation:</u>

   Assume Bla enzymes are instantaneously released by lysing *Klebsiella* when the antibiotic first hits ($t$ = 0). This creates an instantaneous point source of the extracellular Bla concentration, *B*. The spatiotemporal dynamics of *B* are then described by a diffusion equation (Fick's second law), under cylindrical symmetry assumption, as follows:

   $$\frac{\partial B}{\partial t} = D_B \frac{1}{r}\frac{\partial}{\partial r}\left(r\frac{\partial B}{\partial r}\right) \qquad \text{(Equation S1)}$$

   where $D_B$ is the diffusivity of Bla. The time-dependent profile of *B* from the instantaneous point source ($r = Q$) of Bla under cylindrical symmetry is

   $$B(r,t) = \frac{B_M}{4\pi D_B t} e^{\frac{-(r-Q)^2}{4D_B t}}, \qquad r \geq Q > 0$$

   where $B_M$ is the total Bla mass released. $L_t = \sqrt{4D_B t}$ would define a time-dependent diffusion length-scale of Bla's spread. For $D_B$ = 0.15 mm$^2$/h (Supplementary Table 2) and $t$ = 48 h (the main analysis time point for simulations), $L_t$ = 5.37 mm. $L_t$ is ~ 9-fold smaller than the half a domain length (*S* = ~ 45 mm, which is also the dish inner radius for the experiments), suggesting a local Bla effect at our main analysis time points for both the simulations and the experiments (24 h).

   b. <u>Sufficiently fast Bla degradation</u>:

   If Bla gets degraded at a nonnegligible rate, the spatiotemporal dynamics of *B* are then described by a reaction-diffusion equation, under cylindrical symmetry assumption, as follows:

   $$\frac{\partial B}{\partial t} = D_B \frac{1}{r}\frac{\partial}{\partial r}\left(r\frac{\partial B}{\partial r}\right) - d_B B \qquad \text{(Equation S2)}$$

   where $d_B$ is the spontaneous degradation rate constant of Bla.

$\tau = \frac{1}{d_B}$ describes the characteristic timescale for Bla degradation. $L = \sqrt{\frac{D_B}{d_B}}$ describes the characteristic diffusion length scale for Bla spread. There is a vanishingly small probability that Bla molecules travel much farther than the length scale $L$ or survive much longer than $\tau$.

For $d_B$ = 0.02 1/h (Supplementary Table 2), $\tau = \frac{1}{d_B} = 50\,\text{h}$, which is comparable to our main analysis time points for experiments (24 h) and simulations (48 h). With $D_B$ = 0.15 $mm^2$/h (Supplementary Table 2), $L \cong$ 2.74 mm, which is ~16-fold smaller than relevant domain scale (*S* = ~ 45 mm). These suggest a local Bla effect with an approximately steady-state profile ($\frac{\partial B}{\partial t} \cong 0$) at our main analysis time points, i.e.,

$$\frac{\partial^2 B}{\partial r^2} + \frac{1}{r}\frac{\partial B}{\partial r} - \frac{1}{L^2}B = 0 \qquad \text{(Equation S3)}$$

**2. 'Clear zone' radius (*R*)**

Equation S3 is a specific form of Bessel's equation shown below:

$$\frac{d^2 y}{dx^2} + (d-1)\frac{1}{x}\frac{\partial y}{\partial x} - \left(\vartheta - \frac{\mu}{x^2}\right)y = 0$$

The general solution of Bessel's equation is

$$y(x) = C_1 K_\mu\left(x\sqrt{\vartheta}\right) + C_2 I_\mu\left(x\sqrt{\vartheta}\right)$$

where $I_\mu$ and $K_\mu$ are the modified Bessel functions of the first and the second kind, which monotonically grows and decays, respectively, with increasing their arguments[6].

($d = 2$, $\mu = 0$, and $\vartheta = {}^{1}/_{L^2}$ for $y \to B$ and $x \to r$) recovers the Equation S3 above. Thus, its general solution is

$$B(r) = C_1 K_0\left(\frac{r}{L}\right) + C_2 I_0\left(\frac{r}{L}\right)$$

Since Bla effect is local (both $L_t$ and $L$ above are substantially smaller than the relevant domain scale *S*), $\frac{S}{L} \gg 1$

$$B(r = S) = C_1 K_0\left(\frac{S}{L}\right) + C_2 I_0\left(\frac{S}{L}\right) = 0$$

$K_0\left(\frac{S}{L} \gg 1\right) \propto e^{-\infty} \cong 0$. Because $I_0\left(\frac{S}{L} \gg 1\right)$ remains large and positive, the only way to satisfy the above condition is $C_2 = 0$.

Defining $B^*$ as the steady-state Bla enzyme concentration in the *Klebsiella* seeding zone, which is defined by the radius $Q$

$$B(r = Q) = C_1 K_0\left(\frac{Q}{L}\right) = B^*$$

Thus, $C_1 = \frac{B^*}{K_0\left(\frac{Q}{L}\right)}$, and

$$B(r) \cong \frac{B^*}{K_0\left(\frac{Q}{L}\right)} K_0\left(\frac{r}{L}\right), \text{ for } r \geq Q > 0 \quad \text{(Equation S4)}$$

According to Equation S4, *B* decays with increasing $r$ (i.e., moving away from the *Klebsiella* population). Assuming that antibiotic concentration is immediately reduced to zero by $B \geq B_{th}$. Then, the sink radius, $R$, can be defined as the distance at which $B = B_{th}$. Using the latter as a boundary condition for Equation S4,

$$B(r = R) \cong \frac{B^*}{K_0\left(\frac{Q}{L}\right)} K_0\left(\frac{R}{L}\right) = B_{th} \quad \text{(Equation S5)}$$

In our system, $0 < Q < R < S = 45$ mm. That is $0 < \frac{Q}{L} < \frac{R}{L} < 17$ mm. So, the argument of $K_0\left(\frac{Q}{L}\right)$ and $K_0\left(\frac{R}{L}\right)$ can take small to moderate values.

For a moderately valued argument $x$, we approximate $K_0(x)$ by an exponential form with a pre-factor:

$$K_0(x) \approx \sqrt{\frac{\pi}{2(x+1)}} e^{-x} \quad \text{(Equation S6)}$$

After substituting Equation S6 into Equation S5, and doing a Taylor series expansion around $\frac{R}{L} = \frac{Q}{L}$ , we obtain

$$R \cong Q + L\frac{2\ln\left(\frac{B^*}{B_{th}}\right)\left(\frac{Q}{L} + 1\right)}{2\frac{Q}{L} + 3}$$

Because $B^* = B_{th}$ would lead to $\ln\left(\frac{B^*}{B_{th}}\right) = \ln(1) = 0$, we add a manual correction as $\ln\left(\frac{B^*}{B_{th}}\right) \to \ln\left(1 + \frac{B^*}{B_{th}}\right)$. Thus, finally,

$$R \cong Q + L\frac{2\ln\left(1+\frac{B^*}{B_{th}}\right)\left(\frac{Q}{L}+1\right)}{2\frac{Q}{L}+3} \qquad \text{(Equation S7)}$$

#### 3. Spatial profile of the effective *Pseudomonas* spatial expansion ability (*M*)

With its locality, the Bla released by a *Klebsiella* population effectively creates a 'sink' which immediately destroys the antibiotic molecules upon contact. Assuming that the growth media serves as an infinite reservoir of the antibiotic, we have an "infinite source-sink" problem (Fig. 2a, top)[7]. The spatiotemporal dynamics of antibiotic concentration ($A$) is then described by Fick's diffusion equation under cylindrical symmetry,

$$\frac{\partial A}{\partial t} = D_A \frac{1}{r}\frac{\partial}{\partial r}\left(r\frac{\partial A}{\partial r}\right)$$

This problem yields a steady-state $A$ profile, which decreases in the direction from the domain boundary to the 'sink'. At steady-state ($\frac{\partial A}{\partial t} = 0$),

$$D_A \frac{1}{r}\frac{\partial}{\partial r}\left(r\frac{\partial A}{\partial r}\right) = 0$$

This mean $r\frac{\partial A}{\partial r} = C_1$ is a constant. The general solution for $A$ is, thus,

$$A(r) = C_1 \ln r + C_2$$

The ultimate solution must satisfy two boundary conditions as we name them lower and upper boundaries. The lower boundary condition is that the antibiotic concentration (*A*) is zero at 'sink' boundary within which there is a 'clear zone' free of antibiotics:

$$A(r = R) = 0 = C_1 \ln R + C_2$$

$$C_2 = -C_1 \ln R$$

The upper boundary condition is the one that satisfies the maximum (i.e., the initially supplied) concentration of the antibiotic ($A_0$) at a distance sufficiently far away from the 'sink'. If this occurs at $r = S$, which is the experimental domain boundary,

$$A(r = S) = A_0 = C_1 \ln S - C_1 \ln R = C_1 \ln\left(\frac{S}{R}\right)$$

$$C_1 = \frac{A_0}{\ln\left(\frac{S}{R}\right)}$$

Thus,

$$A(r) = A_0 \frac{\ln\left(\frac{r}{R}\right)}{\ln\left(\frac{S}{R}\right)}, \;\; r \,\epsilon\, [R, S] \qquad \text{(Equation S8)}$$

We then introduced a *Pseudomonas* population into the consideration and empirically described the dependence of *Pseudomonas*' effective spatial expansion ability on *A* by a Hill inhibition function based on experimental measurements (Supplementary Fig. 9),

$$M(A) = M_0 \left(\frac{A_C^h}{A_C^h + A^h}\right) \qquad \text{(Equation S9)}$$

where $M(A)$ describes *Pseudomonas*' effective spatial expansion ability as a function of *A*, $M_0$ is the maximum *Pseudomonas*' effective spatial expansion ability under a given condition. Here, $A_C > 0$ and $h > 0$ describe the $A$ value for the half-maximum inhibition and the steepness of the inhibition, respectively, of *Pseudomonas*' effective spatial expansion ability.

Substituting Equation S9 into Equation S8 yields the *Pseudomonas*' effective spatial expansion ability as a function of its distance to the 'sink' ($r \rightarrow d$).

$$M(d) = \frac{M_0}{1+\left(\frac{A_0}{A_C}\right)^h \left(\frac{\ln\left(\frac{d}{R}\right)}{\ln\left(\frac{S}{R}\right)}\right)^h} \qquad \text{(Equation S10)}$$

where $R > 0, S \geq R$, $d \in [R, S]$, $A_0 \geq 0$, and $A_C > 0$. Note that Equation 1 in the main text is the version of Equation S10 for nondimensionalized effective spatial expansion ability and initial antibiotic concentration ($\frac{M}{M_0} \rightarrow M \in [0,1]$ and $\frac{A_O}{A_C} \rightarrow a_0 \in [0\,,\infty]$).

Notice that for sufficiently high values of $h$ and $A_0$, there is a $d = L_K$ at which $M$ abruptly switches between approximately its maximum and minimum values (Fig. 2a, bottom). Thus, $L_K$ has utility as the spatial scale of the keystone engineering.

7 Berg, H. C. *Random Walks in Biology*

*New and Expanded Edition*. REV - Revised edn, (Princeton University Press, 1993).
